# Type 2 Inflammation Drives an Airway Basal Stem Cell Program Through Insulin Receptor Substrate Signaling

**DOI:** 10.1101/2022.10.13.512129

**Authors:** Xin Wang, Nils R. Hallen, Minkyu Lee, Sachin Samuchiwal, Qihua Ye, Kathleen M. Buchheit, Alice Z. Maxfield, Rachel E. Roditi, Regan W. Bergmark, Neil Bhattacharyya, Tessa Ryan, Deb Gakpo, Soumya Raychaudhuri, Dan Dwyer, Tanya M. Laidlaw, Joshua A. Boyce, Maria Gutierrez-Arcelus, Nora A. Barrett

**Author notes:** Corresponding author. Nora A. Barrett, Hale Building for Transformative Medicine, Room 05002R, 60 Fenwood Road, Boston, MA 02115.

## Abstract

**Background:** Chronic rhinosinusitis with nasal polyposis (CRSwNP) is a type 2 (T2) inflammatory disease associated with an increased number of airway basal epithelial cells (BCs). Recent studies have identified transcriptionally distinct BCs, but functional data are lacking and the molecular pathways that support or inhibit human BC proliferation and differentiation are largely unknown.

**Objective:** To determine the role of T2 cytokines in regulating airway BCs

**Methods:** Single cell and bulk RNA-sequencing of sinus and lung airway epithelial cells was analyzed. Human sinus BCs were stimulated with IL-4 and IL-13 in the presence and absence of IL4R inhibitors. Confocal analysis of human sinus tissue and murine airway was performed. Murine BC subsets were sorted for RNA sequencing and functional assays. Fate labeling was performed in a murine model of tracheal injury and repair.

**Results:** Here we find two subsets of BCs in human and murine respiratory mucosa distinguished by the expression of BC adhesion molecule (BCAM). BCAM expression identifies airway stem cells among P63+KRT5+NGFR+ BCs. In the sinonasal mucosa, BCAM^hi^ BCs expressing *TSLP*, *IL33*, *CCL26,* and the canonical BC transcription factor *TP63* are increased in patients with CRSwNP. In cultured BCs, IL-4/13 increases expression of *BCAM* and *TP63* through an Insulin Receptor Substrate (IRS)-dependent signaling pathway that is increased in CRSwNP.

**Conclusions:** These findings establish BCAM as a marker of airway stem cells among the BC pool and demonstrate that airway epithelial remodeling in T2 inflammation extends beyond goblet cell metaplasia to the support of a BC stem state poised to perpetuate inflammation.

**CAPSULE SUMMARY:** Type 2 cytokines drive an airway stem cell program through IRS signaling

**KEY MESSAGES:**

- Two subsets of airway BCs have distinct transcriptional signatures and function
- High levels of BCAM expression mark the earliest BC progenitor
- IL-4 and IL-13 upregulate BCAM and P63 in an IRS-dependent fashion which prevents BC differentiation to secretory epithelial cells
- BCAM^hi^ BCs are increased in CRSwNP

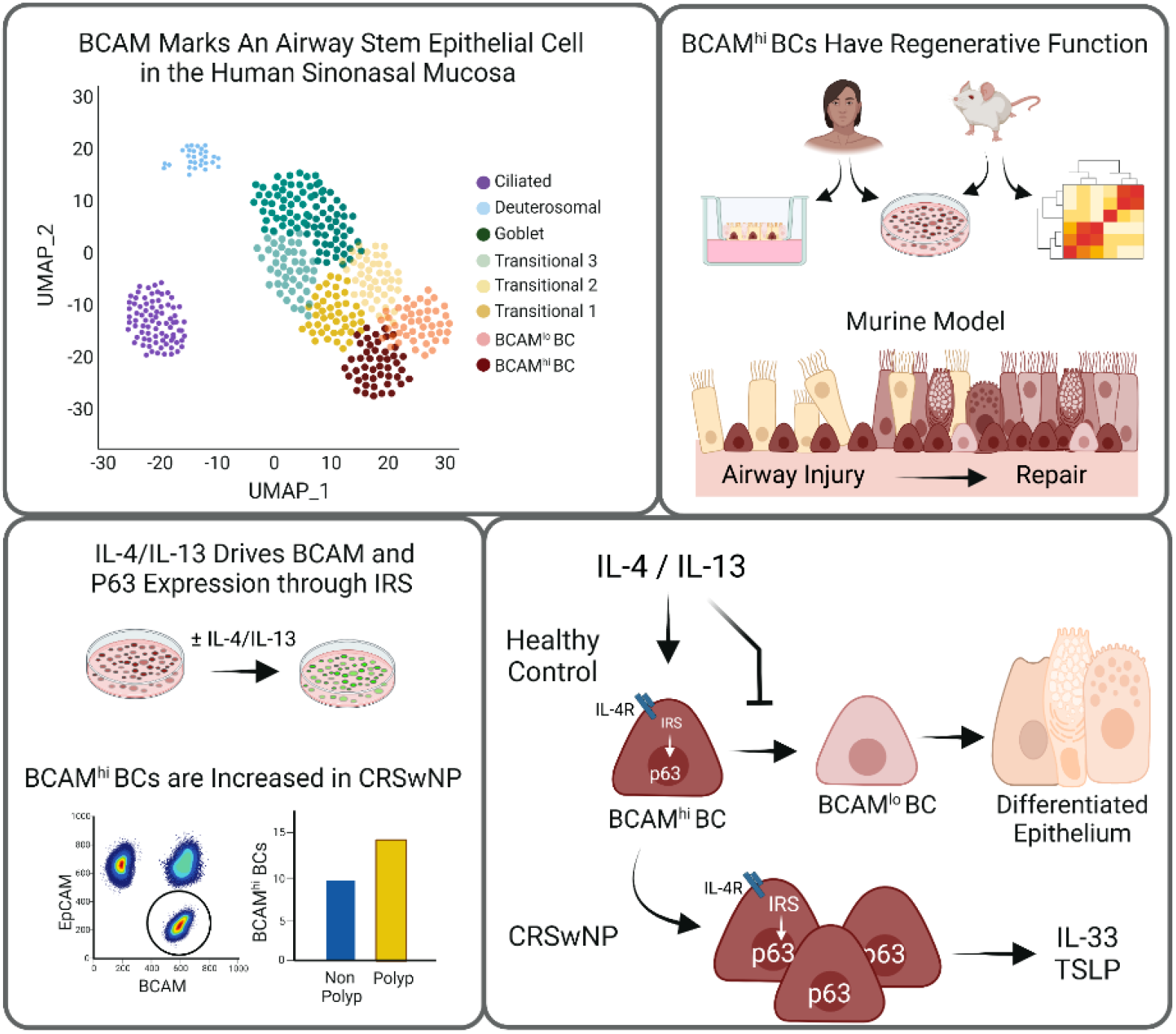

## INTRODUCTION

Tissue-resident stem cells exhibit remarkable plasticity, responding to local damage by regenerating diverse differentiated cell types. This plasticity ensures their ability to restore homeostasis after injury but also endows them with the capacity to remodel the tissue microenvironment and adapt to tissue stress. A canonical example of tissue remodeling is intestinal goblet cell metaplasia, which plays an essential role in host defense against helminths. Here local immunocytes generate interleukin-13 (IL-13) which drives the differentiation of epithelial cells (EpCs) into goblet cells that secrete mucus and thereby facilitate helminth expulsion (1–3). While appropriate tissue adaptation requires that stem cells integrate local environmental cues from diverse sources including niche mesenchymal cells, tissue immunocytes, and even their own progeny, in most circumstances the signals that mediate remodeling and the features of remodeling that are beneficial or detrimental to the host are poorly understood.

In the respiratory tract, studies using single cell RNA-sequencing (scRNA-seq) and immunofluorescence have identified variations in the abundance of EpC subsets in distinct disease states. This includes an expansion of ciliated cells in cystic fibrosis (4); an increase in IL-25-secreting tuft cells in allergic fungal rhinosinusitis and chronic rhinosinusitis with nasal polyposis (CRSwNP) (5–7); and an increase in neuroendocrine cells in diffuse idiopathic pulmonary neuroendocrine cell hyperplasia (8–10), asthma (11), and neuroendocrine cell hyperplasia of infancy (12). These specialized EpCs derive from basal cell (BC) progenitors, suggesting that alterations in BC programs likely account for these variations. Indeed, scRNA-seq has detected transcriptionally distinct BC subsets in the respiratory tract (13–15), but functional annotation of these cell types is lacking.

In Western Countries, CRSwNP is a type 2 (T2) inflammatory disease of the sinonasal airway which is often associated with asthma. Patients with this disorder have eosinophilia in the airways and peripheral blood and respond to treatment with monoclonal antibody blockade of IL4Rα, a component of both the type I and type II IL-4 receptors that bind IL-4 and IL-13 (16). We previously reported that BCs from patients with CRSwNP accumulated in the sinonasal mucosa and failed to differentiate normally, but the mechanism by which this occurs is unknown.

Here we report two subsets of KRT5+ BCs in the human sinonasal mucosa distinguished by expression of BC adhesion molecule (BCAM). As compared to BCAM^lo^ BCs, BCAM^hi^ BCs express higher levels of the canonical stem marker *TP63* and have the capacity to support both ciliated and secretory lineages in *ex vivo* air-liquid interface (ALI) culture, reflecting their stem function. Trajectory analysis of human samples and fate tracking experiments in mice confirm that BCAM^hi^ BCs are the progenitors of BCAM^lo^ BCs. In human sinonasal mucosa, BCAM^hi^ BCs are increased in number in CRSwNP, as compared to controls, and expression of both *BCAM* and *TP63* correlates with epithelial genes induced by T2 cytokines, suggesting that T2 cytokines may maintain stem function. Ex-vivo BC culture demonstrates that IL-4 and IL-13 reinforce expression of BCAM and P63 through an IRS-dependent signaling pathway that is overexpressed in CRSwNP. These results demonstrate that functionally distinct airway progenitor cells are distinguished by expression of BCAM. Additionally, they demonstrate that the T2 high environment of CRSwNP plays a profound role in remodeling the airway epithelial BC compartment to drive a stem cell with potent pro-inflammatory capacity.

## RESULTS

### BCAM marks a multipotent progenitor cell among KRT5^+^NGFR^+^ITGA6^+^ BCs in the human respiratory mucosa

We previously reported the impaired differentiation of airway EpCs in patients with CRSwNP (13). BC dysplasia was associated with increased BC number, upregulated Wnt signaling, profound chromatin remodeling, and an altered expression of key transcription factors. To better delineate EpC differentiation, we reanalyzed our scRNA-seq dataset from sinus surgeries (13). Using *Harmony* (17), we integrated data across donors and patient subtypes (CRSsNP and CRSwNP) in principal component (PC) space, reclustered non-proliferating surface airway secretory and ciliated EpCs (**Fig. S1A-D**), and assessed established EpC markers (**Table 1**) to define common EpC states across diseases (**Fig. S1E**). Here we identified two BC subsets present in the sinonasal tissue of patients with both CRSwNP and CRSsNP (**Fig. 1A-C**). One subset (cluster 0) expressed high levels of the BC markers *KRT5, KRT15,* and *TP63* (18–20); the stem marker *CD44* (21); and the cell surface receptor *BCAM* (FDR < .05, **Fig. 1B-C, Table S4**). The second BC subset (cluster 1) expressed higher levels of the oncogene *MALAT1* (22) and many ribosomal genes, suggesting ribosomal biogenesis. Differentiating transitional cells (clusters 2-4) were marked by increasing expression of the Notch pathway gene *HES1* (23), the luminal BC marker *KRT8* (24), and the club cell markers *SCGB1A1* and *TFF3* (25, 26); while mature secretory cells (cluster 5) were marked by *MUC5AC* (14); deuterosomal cells (cluster 6) were marked by *CDC20B*, *PLK4*, and *FOXJ1* (27); and ciliated cells (cluster 7) were marked by *PIFO*, *CAPS*, and *FOXJ1* (15). Cluster 5 was composed of both goblet and club secretory cells expressing *MUC5AC* or *MUC5B* and *SCGB1A1*, respectively (**Fig. S1F**).

**Table 1.** Established Markers of Airway Epithelial Cells.

| **Table 1. Established Markers of Airway Epithelial Cells** |  |  |
| --- | --- | --- |
| **Cell Type** | **Marker** |  |
| **BCs** | NGFR | Nerve growth factor receptor |
|  | GSIB4* | Lectin Griffonia Simplicifolia-IB4 |
|  | ITGA6 | Integrin subunit alpha 6 |
|  | KRT5 | Keratin 5 |
|  | TP63 | Tumor protein 63 |
|  | CD44 | CD44 molecule |
|  | PDPN | Podoplanin |
| **Luminal Progenitor BCs** | KRT8 | Keratin 8 |
| **Secretory Club Cells** | SCGB1A1 | Secretoglobulin family 1A member 1 |
| **Goblet Cells** | MUC5AC | Mucin 5AC |
| **Ciliated Cells** | FOXJ1 | Forkhead box J1 |
| | Acetyl-α<br>Tubulin | Acetylated alpha tubulin |
| *GSIB4 lectin is a marker of murine BCs but not human |  |  |

**Fig. 1.**
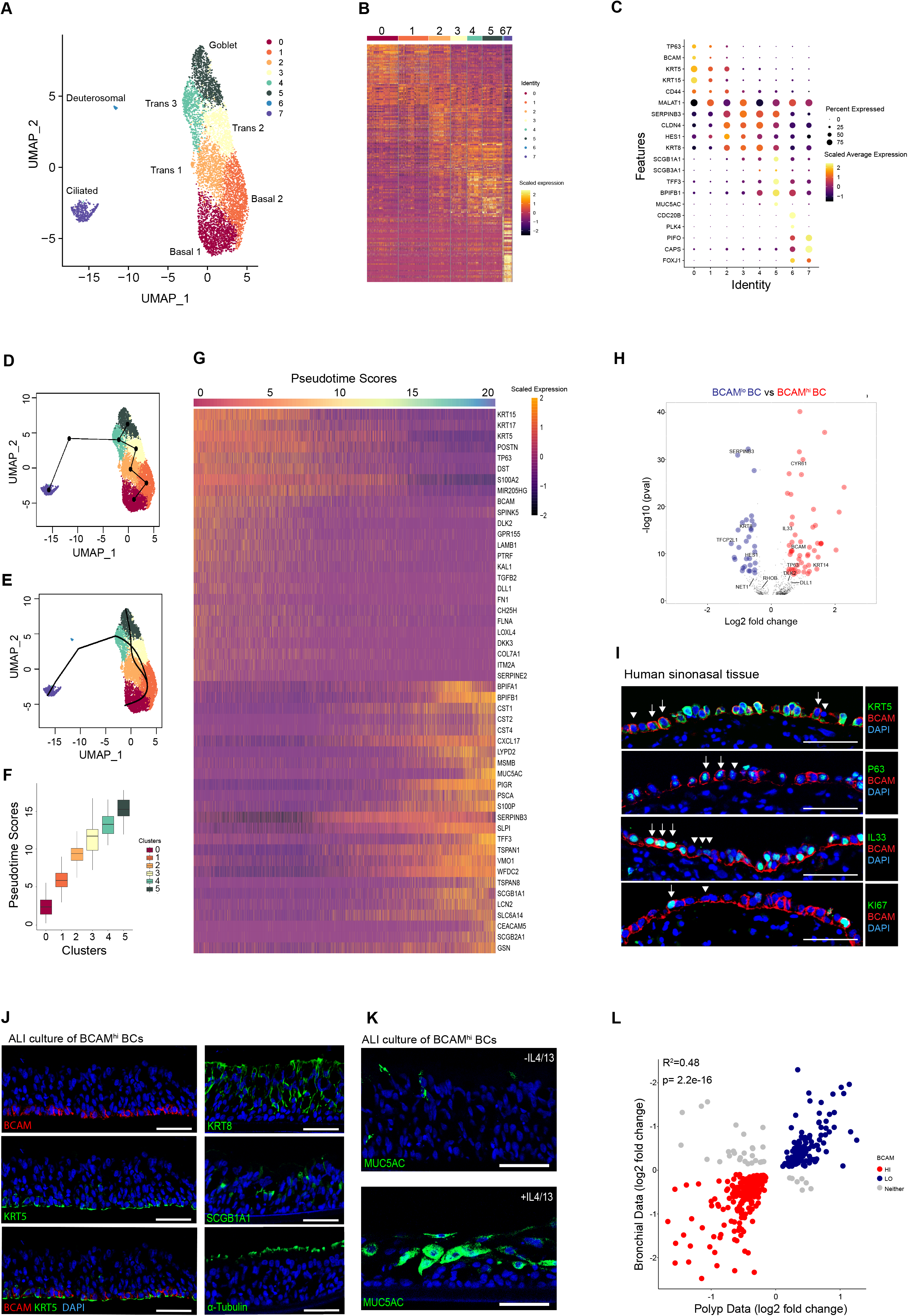
BCAM marks a multipotent progenitor among KRT5^+^NGFR^+^ITGA6^+^ BCs in the human respiratory mucosa. (**A**) UMAP representation of 6970 EpCs from 12 donors. (**B**) Heatmap of scaled gene expression for the top 30 genes identified by Wilcoxon rank sum test and auROC analysis. (**C**) Dot plot of epithelial markers across clusters. (**D**) Cluster-based minimum spanning tree of lineages overlaid on UMAP. (**E**) Cell assignment to smooth principle curves with lineage-specific pseudotimes. (**F**) Plot of pseudotime scores from cells assigned to the secretory lineage. (**G**) Heatmap of scaled gene expression for the topmost significant genes associated with changes in pseudotime (Bonferroni corrected p-values). (**H**) Volcano plot of log_2_fold-change between Basal 0 (BCAM^hi^) BCs and Basal 1 (BCAM^lo^) BCs. Significant genes with increased expression in BCAM^hi^ (red) and BCAM^lo^ (blue) (P < 1.5e-05, Bonferroni threshold). (**I**) Representative immunostaining of human sinonasal tissue from a patient with CRSwNP. Arrow indicates BCAM^hi^ BCs, arrowhead indicates BCAM^lo^ BCs. The scale bar represents 50 μm. n = 3. (**J**) Representative immunostaining on ALI cultures derived from BCAM^hi^ BCs. The scale bar represents 50 μm. n = 3. (**K**) Representative immunostaining on ALI cultures derived from BCAM^hi^ BCs treated with/without IL4/13. n = 3. (**L**) Coefficient of determination of log_2_fold changes of shared genes from sinonasal BCs (BCAM^hi^ vs BCAM^lo^) and bronchial BCs (BC1 vs BC2).

Trajectory analysis of these clusters suggested two distinct trajectories with a serial progression from *TP63^hi^* BC progenitors (cluster 0) to *MALAT1*^+^ BCs (cluster 1), *KRT8*^+^ *HES1*^hi^ transitional cells (clusters 2-4), and then to either fully differentiated MUC5AC^+^ goblet cells (cluster 5) or to *PIFO*^+^ ciliated EpCs (cluster 7) (**Fig. 1D-E**). Accordingly, cells in the secretory trajectory from clusters 0-5 (**Fig. S1G**) demonstrated progressively increasing pseudotime scores (**Fig. 1F**). Using generalized linear modeling across pseudotime for EpCs in the secretory trajectory, we demonstrated 2278 genes that were either highly positively or negatively associated with the pseudotime score (P < 4.23e-06, Bonferroni threshold, **Fig. 1G, Tables S5-6**). We detected increasing expression of canonical secretory genes (*MUC5AC, SERPINB3*, *SLPI*, *TFF3, SCGB1A1, CEACAM5*) across pseudotime and loss of basal cell markers (*KRT5*, *KRT15*, *TP63*), as expected. Additionally, we saw that expression of the cluster 0 BC marker *BCAM* (**Fig. 1C**) was inversely associated with differentiation, suggesting that it’s expression may distinguish BC states.

Direct comparison of cluster 0 BCAM^hi^ BCs and cluster 1 BCAM^lo^ BCs demonstrated 142 genes that were differentially expressed between these clusters (P < 1.5e-05, Bonferroni threshold, **Fig. 1H, Table S7**). BCAM^hi^ BCs expressed higher levels of many genes associated with epithelial progenitor cells. This included the Yap target gene *CYR61/CCN1* (28); WNT target genes *MMP10* (29) and *MYC* (30); the NOTCH ligand *DLL1* (31) and the NOTCH inhibitor *DLK2* (32); the anti-apoptotic genes *BIRC3* (33) and *EMP1* (34, 35); the proliferation-associated gene *ZFP36L2* (36); and transcription factors required for stem cell maintenance such as *TCF12* (37) and *TP63* (19, 20, 38). BCAM^hi^ BCs also expressed higher levels of the T2 cytokine and transcriptional repressor *IL33* (39, 40) and showed a trend to increased *TSLP* that was not significant after Bonferonni adjustment. Additionally, BCAM^hi^ BCs expressed higher levels of genes encoding growth factors and extracellular matrix components such as *POSTN* (periostin), *FN1* (fibronectin), *CTGF* (connective tissue growth factor), *LAMB1* (laminin subunit B1), and *LAMB3* (laminin subunit B3). By contrast, BCAM^lo^ BCs expressed higher levels of genes associated with differentiation including *SERPINB3* (41)*, HES1* (23)*, KRT8* (31) and the ETS transcription factor *ELF3* (42, 43), suggesting that increased *BCAM* expression identifies early airway progenitors among the BC pool.

Confocal images of the epithelium in CRSwNP confirmed variable expression of BCAM with some KRT5^+^ BCs expressing BCAM solely on the basal surface (designated BCAM^lo^) and other KRT5^+^ BCs expressing BCAM circumferentially (designated BCAM^hi^)(**Fig. 1I, top row**). BCAM^hi^ BCs expressed p63 (**Fig. 1I, second row**), IL-33 (**Fig. 1I, third row**), and Ki67 (**Fig. 1l, fourth row**) but none of these proteins were *exclusively* expressed in BCAM^hi^ BCs. Flow cytometry also demonstrated variable BCAM expression among lin^−^(CD45^−^CD90^−^CD31^−^) EpCAM^lo^NGFR^hi^ BCs (**Fig. S1H**). Remarkably, both BCAM^hi^ and BCAM^lo^ BCs had similar staining for integrin alpha 6 (18, 44), and little distinction in the expression levels of the BC markers podoplanin (31) and nerve growth factor receptor (NGFR) (18) (**Fig. 1I**). Sorted BCAM^hi^ BCs from healthy controls and from CRSwNP were passaged and differentiated in ALI cultures that supported the growth of KRT5^+^ BCs, KRT8^+^ luminal cells, SCGB1A1^+^ club cells, acetylated tubulin^+^ ciliated cells, and Muc5AC^+^ goblet cells (**Fig. 1J-K**). By contrast, EpCAM^lo^NGFR^hi^BCAM^lo^ BCs survived only limited passages *ex vivo* and could not support differentiation in ALI cultures. Taken together, these results suggested that BCAM expression distinguishes sinonasal stem cells among BCs.

Notably, examination of a published scRNA-seq dataset from bronchial mucosa (14) revealed that *BCAM* also marks a subset of bronchial BCs from healthy donors (**Fig. S1J**) with co-expression of *TP63* and *IL33.* After assessment of differential gene expression between BCAM^hi^ and BCAM^lo^ bronchial BCs, we analyzed differentially expressed genes that were detected in both the bronchial and polyp BC subsets. This demonstrated a strong correlation in gene programs across datasets (**Fig. 1L, Table S8**), indicating that a similar BCAM^hi^ BC population exists in the lower airway.

### BCAM marks an airway stem cell among KRT5^+^NGFR^+^ITGA6^+^P63^+^ BCs in the murine trachea

The limited progenitor capacity of BCAM^lo^ BCs in *ex vivo* culture was striking, as even differentiated club cells are reported to retain some progenitor capacity in the appropriate context (45, 46). Thus, we next sought to assess these populations *in vivo*, and turned to the murine airway. We first established the presence of two identifiable BC subsets. In the sinonasal mucosa, confocal microscopy demonstrated some basal epithelial cells that expressed high levels of BCAM and P63, and other basal epithelial cells that expressed low levels of BCAM and lacked P63 (**Fig. S2A**). However, detailed assessments were limited by prolonged protocols to decalcify the tissue before staining (47, 48), thus we turned to the trachea. Here again we identified two patterns of BCAM staining on KRT5^+^ BCs, with circumferential expression of BCAM on some KRT5^+^ BCs and focal basolateral expression of BCAM on other KRT5^+^ BCs (**Fig. 2A**). P63 (**Fig. 2A**) was primarily expressed in BCAM^hi^ BCs, consistent with a BC progenitor. As isolation and analysis of tracheal BCs was readily performed, we established a flow cytometric panel for their evaluation in naïve mice. Within the conventional BC gate (lin^−^EpCAM^lo^GSIB4^hi^) (49, 50), we again identified two EpC subsets distinguished by BCAM expression (**Fig. 2B, Fig. S2B**). There was no distinction in the level of EpCAM expression between BCAM^hi^ and BCAM^lo^ BCs (**Fig. 2C**), and both populations expressed high levels of the BC markers KRT5, P63, ITGA6 and NGFR (**Fig. 2D**). Notably, KI67+ cells were dominantly detected in the BCAM^hi^ group, which was confirmed by confocal microscopy (**Fig. 2E**). By contrast, the GSIB4^−^BCAM^−^ EpCs lacked expression of each BC marker, but expressed higher levels of molecules marking differentiated EpCs including EpCAM, KRT8, SCGB1A1, SSEA1, and choline acetyl transferase (**Fig. S2C-E**).

**Fig. 2.**
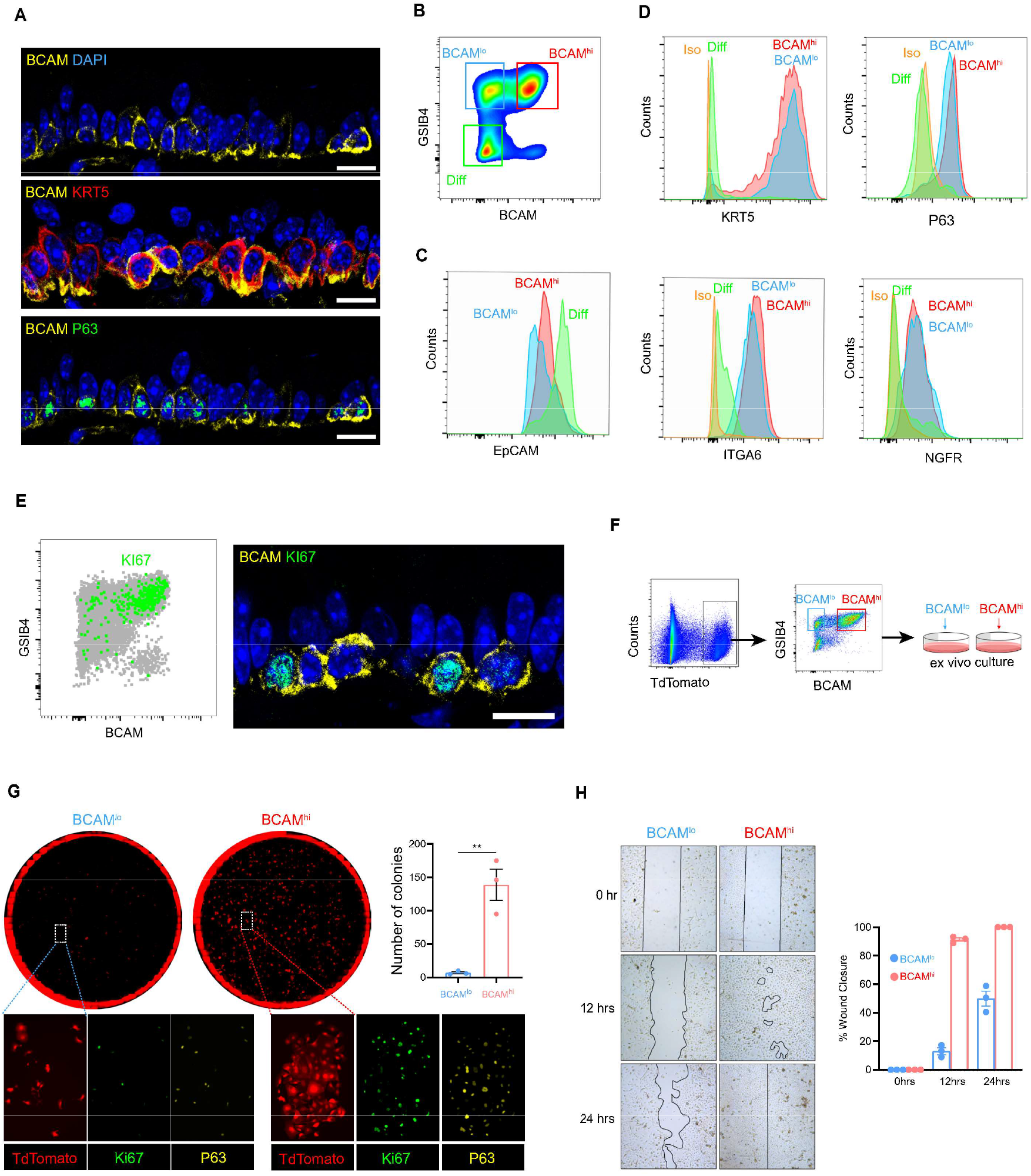
BCAM marks an airway basal stem cell among KRT5^+^NGFR^+^ITGA6^+^P63^+^ BCs in the naive murine trachea. (**A**) Representative immunostaining in naïve murine trachea. The scale bar represents 10 μm. (**B**) Flow cytometric panel of BCAM^hi^ BCs, BCAM^lo^ BCs and differentiated EpCs (Diff). (**C**) Expression of EpCAM in each group. (**D**) Expression of canonical BC markers in each group. (**E**) KI67 expression in naïve tracheal airway epithelium assessed by flow cytometric staining (left) and confocal microscopy (right). The scale bar represents 10 μm. (**F**) Schema depicting the isolation and *ex vivo* culture of BCAM^hi^ BCs and BCAM^lo^ BCs from KRT5^CrerERT2^: R26^tdTomato^ mice, detailed information in Materials and Methods. (**G**) Colony forming assay on sorted BCAM^hi^ BCs and BCAM^lo^ BCs with immunostaining. Number of colonies were calculated using image J. Data are shown as mean ± SEM (n = 3; **p < 0.01, unpaired two-tailed t test). (**H**) Wound healing assay on sorted BCAM^hi^ BCs and BCAM^lo^ BCs. Images were taken at 0, 12 and 24 hr. The percentage of wound closure was calculated using image J. Data are mean ± SEM (n = 3; p < 0.02, linear regression). All immunostaining is representative of n = 3.

We next used a tamoxifen-inducible KRT5 reporter mouse to sort tracheal BCAM^hi^ and BCAM^lo^ BCs and expand them in submerged culture (**Fig. 2F-G**). BCAM^hi^ BCs formed larger Ki67^+^ and P63^+^ colonies that were not present in BCAM^lo^ BC cultures (**Fig. 2G**). Finally, BCAM^hi^ BC cultures were grown to confluency, scratched to clear cells from the center, and followed for 24 h to assess wound healing (**Fig. 2H**). This demonstrated that BCAM^hi^ BCs were more efficient than BCAM^lo^ BCs at closing a wound. Thus, both classical markers of replication and functional assays for proliferation and wound healing demonstrated that BCAM distinguishes a murine airway stem cell among KRT5^+^ BCs.

### EpCs differentiate from BCAM^hi^ to BCAM^lo^ BCs in a model of airway injury and inflammation

Having established the *ex vivo* behavior of murine BCAM^hi^ and BCAM^lo^ BCs, we next assessed their behavior in a murine model of airway inflammation and injury using the repetitive inhalation of the mold aeroallergen *Alternaria alternata* (ALT) over 1 or 2 weeks (**Fig. 3A**). Hematoxylin and eosin staining demonstrated epithelial injury by day 7 and regeneration by day 14 (**Fig. 3B**). Flow cytometry on day 7 demonstrated a reduction of BCAM^hi^ BCs, but an increase in BCAM^int^ (intermediate) and BCAM^lo^ populations (**Fig. 3C**). By day 14, we found recovery of BCAM^hi^ BCs, a decrease in the BCAM^int^ population, and a further increase in BCAM^lo^ BCs (**Fig. 3C**). Notably, after these repetitive challenges, BCAM^lo^ BCs expressed some secretory markers including SSEA1 and Muc5AC (**Fig. 3D, Fig. S3A**), suggesting early differentiation. Flow cytometry and confocal images showed that while an increasing number of BCAM^lo^ BCs and differentiated EpCs expressed Ki67 over the challenges, BCAM^hi^ BCs were the dominant cell type expressing Ki67 at all timepoints (**Fig. 3E, Fig. S3B-C**). Taken together, these results suggested that BCAM marks a renewable stem progenitor responsive to airway injury, and that BCAM^hi^ BCs may give rise to BCAM^lo^ BCs and then to differentiated EpCs.

**Fig. 3.**
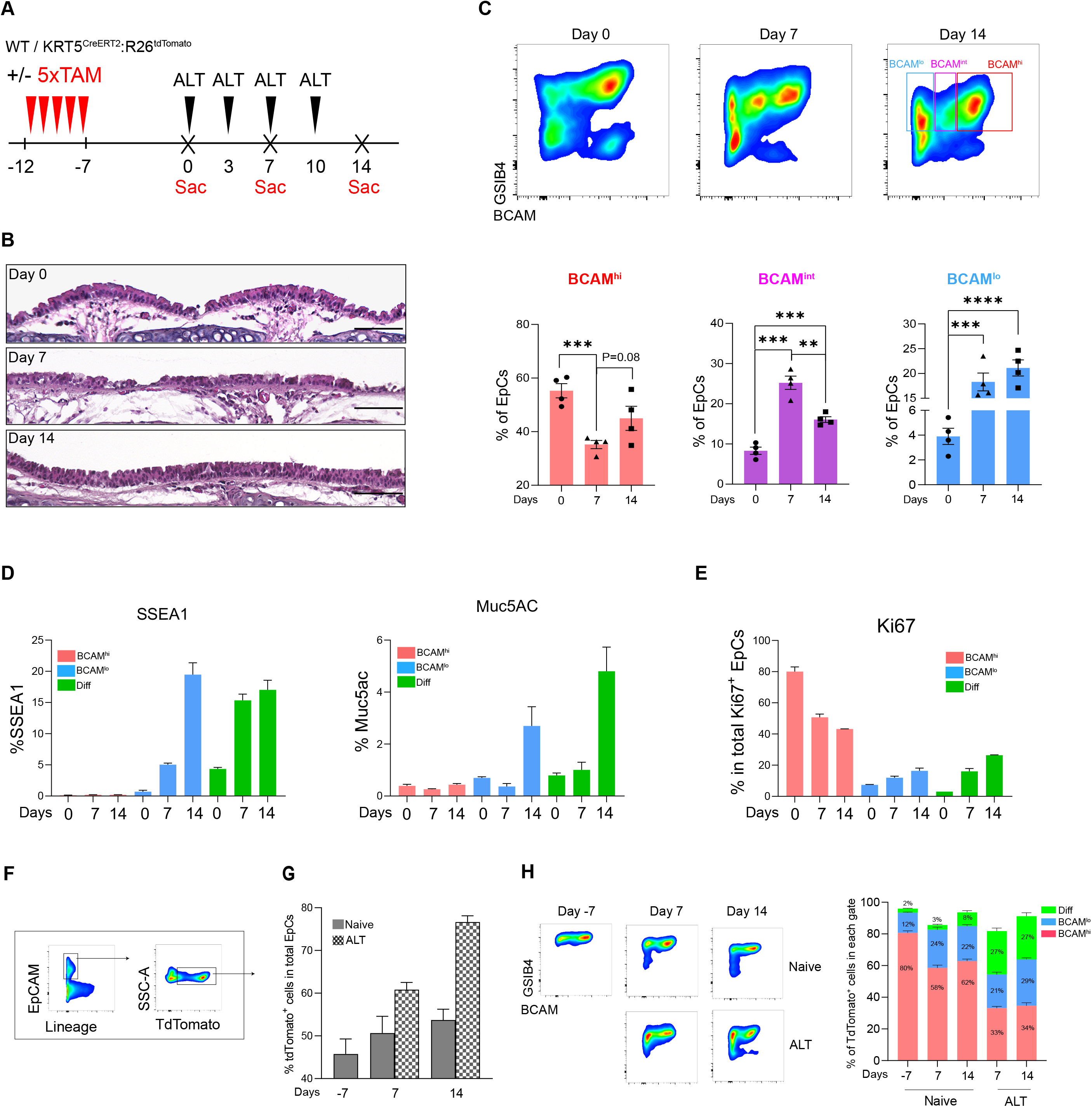
Fate labeling demonstrates a trajectory from BCAM^hi^ BCs to BCAM^lo^ BCs and then to differentiated EpCs. (**A**) Experimental schema. (**B**) H&E staining of mouse trachea. The scale bar represents 100 μm. Staining is representative of n = 3. (**C**) Flow cytometric panel showing the percentage of BCAM^hi^ BCs, BCAM^int^ BCs and BCAM^lo^ BCs in naïve and challenged airways. Data are shown as mean ± SEM (n = 4; **p < 0.01, *** p < 0.001, unpaired two-tailed t test). (**D**) Percentage of SSEA1^+^ cells and Muc5AC^+^ cells in each EpC subset. Data are shown as mean ± SEM (n = 3). (**E**) Percentage of Ki67+ cells in each EpC subset. Data are shown as mean ± SEM (n = 3). (**F**) Gating strategy identifying TdTom+ EpCs in KRT5^CrerERT2^: R26^tdTomato^ mice. (**G**) Percentage of TdTom+ labeled EpCs out of total EpCs at the indicated timepoints. Data are shown as mean ± SEM (n = 3). (**H**) A representative image of GSIB4 and BCAM staining on TdTom+ EpCs (left). Percent of TdTom+ EpCs that fall within each gate in naïve and ALT at the indicated timepoints (right). Data are shown as mean ± SEM (n = 3).

To confirm that BCAM^hi^ BCs are the progenitors of BCAM^lo^ BCs we performed fate mapping using tamoxifen-treated, ALT-challenged KRT5^CreERT2^R26^tdTomato^mice. Mice were treated with 5 doses of tamoxifen (TAM) to label KRT5^+^ BCs, rested for one week, and then treated with intranasal ALT over 1 or 2 weeks (**Fig. 3A**). As expected, the percent of EpCs labelled with tdTomato increased over the ALT challenges (**Fig. 3F-G**). While at day 0 tdTomato labelling was restricted to the basal layer (**Fig. S3D-E**), labelling was detected in SCGB1A1+ and acetylated tubulin+ cells over the ALT protocol. Assessment of lin-EpCAM^+^ TdTomato^+^ cells demonstrated that approximately 80% of labelled cells at day −7 were BCAM^hi^ BCs, only 10-15% of labelled cells were BCAM^lo^ BCs, and < 5% were differentiated EpCs, while some labelled cells fell between these gates (**Fig. 3H**). At days 7 and 14 after airway injury, a higher percentage of labelled cells appeared in the BCAM^lo^ and differentiated EpC gates such that, by day 14, labelled cells appeared nearly equally distributed across BCAM^hi^, BCAM^lo^, and differentiated EpC gates (**Fig. 3H**). Conversely, when gating each EpC population first, we observed rapid labelling of BCAM^hi^ BCs in the naïve trachea, with increasing labelling of the BCAM^lo^ BCs and the differentiated EpCs over 14 days (**Fig. S3F**). These data are consistent with an EpC differentiation trajectory that begins with BCAM^hi^ BCs progresses to BCAM^lo^ BCs and then to differentiated EpCs.

### Transcriptional profile of murine BCAM^hi^ and BCAM^lo^ BCs

As our low resolution scRNA-seq provided only a limited assessment of these BC subsets that displayed such distinct *ex vivo* and *in vivo* behaviors, we next isolated them from naïve murine trachea, and assessed their transcriptional differences with bulk RNA-seq (**Fig. S4A**). Principal component analysis demonstrated that PC1 separated samples by stage of differentiation (**Fig. 4A, Table S9**). Direct comparison of BCAM^hi^ and BCAM^lo^ BCs demonstrated that BCAM^hi^ BCs expressed higher levels of many of the same markers detected at elevated levels in human BCAM^hi^ BCs including *Krt5*, *Trp63*, *Ccn1* (*CYR61*), *Lamb3, Jag2*, *Dlk2*, and *Bcam,* and higher levels of additional BC markers *Ngfr*, *Pdpn*, *Krt14*, and *Krt17* (FDR < .05 **Fig. 4B-C, Fig. S4B, Table S10-11**). *Il33* was poorly detected in each group. BCAM^hi^ and BCAM^lo^ BCs expressed significantly different levels of genes critical for EpC differentiation and maintenance of stemness including genes in the Wnt (**Fig. 4D**), Hippo-Yap (**Fig. 4E**), Notch (**Fig. 4F**), Rho (**Fig. 4G**), and Ras (**Fig. 4H**) pathways. While only a few genes in each of these pathways were recovered in the human scRNA-seq dataset, all but one of them (*CCND1*) were significantly increased in BCAM^hi^ BCs, as compared to BCAM^lo^ BCs (**Fig. S4C**). Transcription factors that were significantly different in the murine data included the regulator of embryonic morphogenesis *Hoxd8* (51); the driver of respiratory and epidermal basal cell proliferation *Vdr* (52, 53); the tumor suppressors *Klf10, Klf11,* and *Trp53* (54–56); and 17 transcripts encoding zinc-finger proteins, all of which were increased in BCAM^hi^ BCs (**Fig. S4D, Table S12**). Among the top transcription factors differentially regulated was *Trp63* (**Fig. S4D**), the murine homologue of *TP63* that was detected in human sinonasal BCAM^hi^ BCs (**Fig. 1C, 1H**). Assessment of a previously reported list of 175 genes with *TP63* binding sites (57) showed that 34 were differentially expressed across BCAM^hi^ and BCAM^lo^ BC subsets (**Fig. 4I**). Moreover, 10 were among the top 250 genes upregulated in BCAM^hi^ BCs, suggesting a potential link between BCAM and *Trp63.* Assessment of our previously published bulk RNA-seq dataset from human sinonasal BCs from CRSwNP demonstrated strong correlation between *BCAM* expression and *TP63* (*R^2^* = 0.85, p = 0.001 **Fig. 4J, Table S13**), suggesting a potential link between *BCAM* and *TP63* expression in human BCs.

**Fig. 4.**
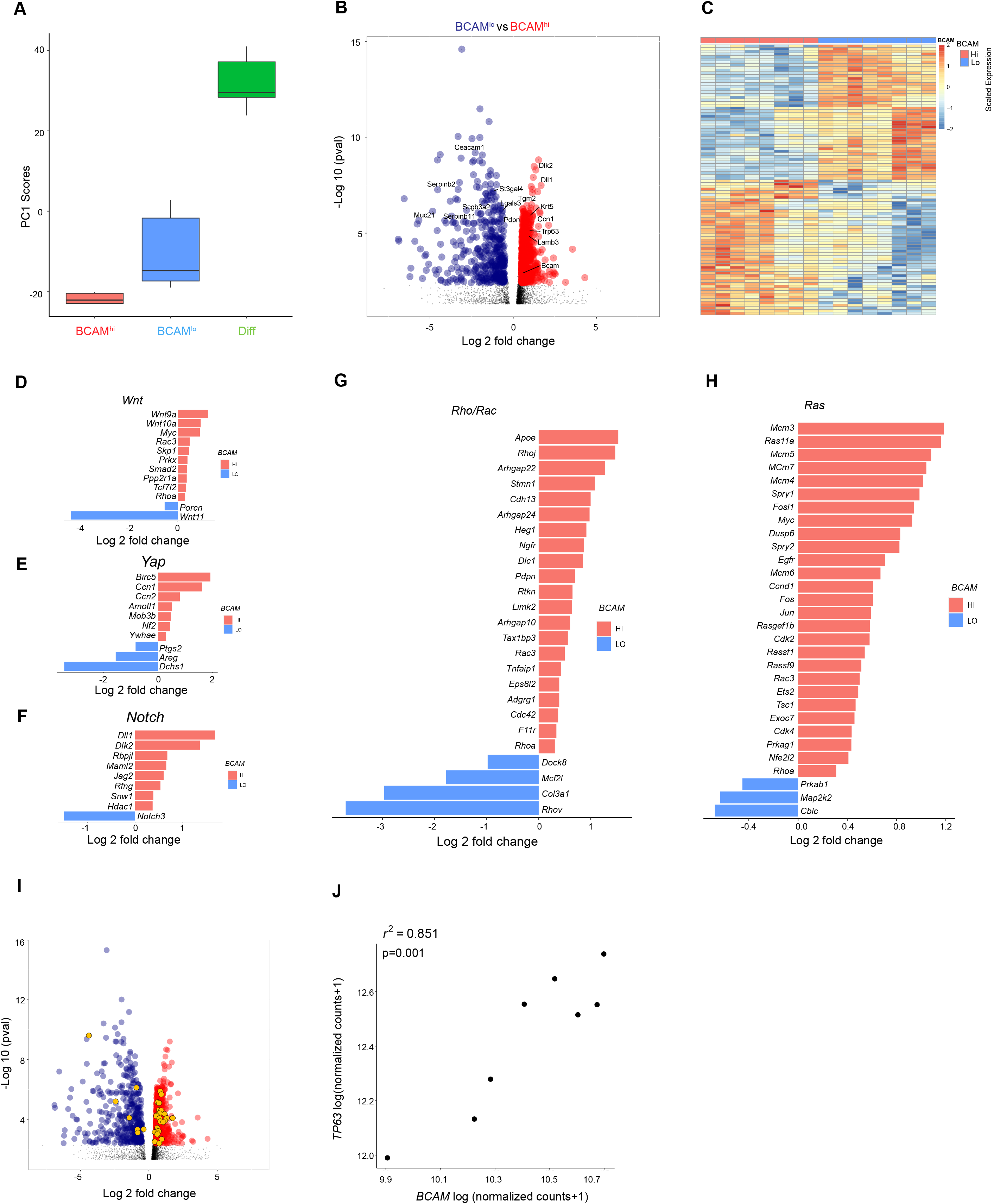
BCAM^hi^ BCs are enriched in canonical stem cell signaling pathways. (**A**) Box plot of principal component 1 scores from BCAM^hi^ BCs, BCAM^lo^ BCs, and Diff EpCs. (**B**) Volcano plot showing differentially expressed genes in BCAM^hi^ BCs compared with BCAM^lo^ BCs. Highlighted genes are significantly enriched in BCAM^hi^ BCs (red) and BCAM^lo^ BCs (dark blue), (Benjamini-Hochberg p-value < .05). (**C**) Heatmap of the top 100 most significant differentially expressed genes in BCAM^hi^ BCs and BCAM^lo^ BCs, (Benjamini-Hochberg p-value < .05). (**D-H**) Significant differentially expressed genes (Benjamini-Hochberg p-value < .05) between BCAM and BCAM BCs associated with *Wnt* signaling (D), *Hippo* signaling (E), *Notch* signaling (F), *Rho/Rock* signaling (G) and *Ras* signaling (H). (**I**) Volcano plot showing differentially expressed genes up in BCAM^hi^ BCs (red) compared with BCAM^lo^ BCs (blue). The significant differentially expressed genes that are P63 target genes are colored in orange. (**J**) Correlation between *BCAM* and *TP63* expression (normalized counts; DESeq2 median of ratios) in polyp BCs bulk RNA-sequencing.

### T2 / IRS-dependent regulation of BCAM

*TP63* was previously reported to be induced by the T2 cytokine IL-13 in human keratinocytes (58), thus we hypothesized that *BCAM* and *TP63* may be similarly regulated in the sinonasal mucosa. First, we assessed *BCAM* and *TP63* expression in bulk RNA-seq of sinonasal BCs from CRSwNP, a T2 high disease, and from non-polyp controls (CRSsNP). Polyp BCs expressed higher levels of *BCAM* and *TP63* and higher levels of canonical T2-inducible epithelial genes such as *ALOX15*, *POSTN*, *IL33*, and *TSLP* (**Fig. 5A, Table S14**). There was a trend to increased expression of *CCL26* that was not significant. Additionally, flow cytometry demonstrated that the percent of BCAM^hi^ BCs among lin^−^EpCAM^lo^NGFR^+^ cells was higher in CRSwNP than CRSsNP (**Fig. 5B**). Interestingly, assessment of TP63-dependent genes also showed an increase in BCs from CRSwNP, as compared to CRSsNP (**Fig. 5C**), suggesting an increase in this stem program in this T2 disease.

**Fig. 5.**
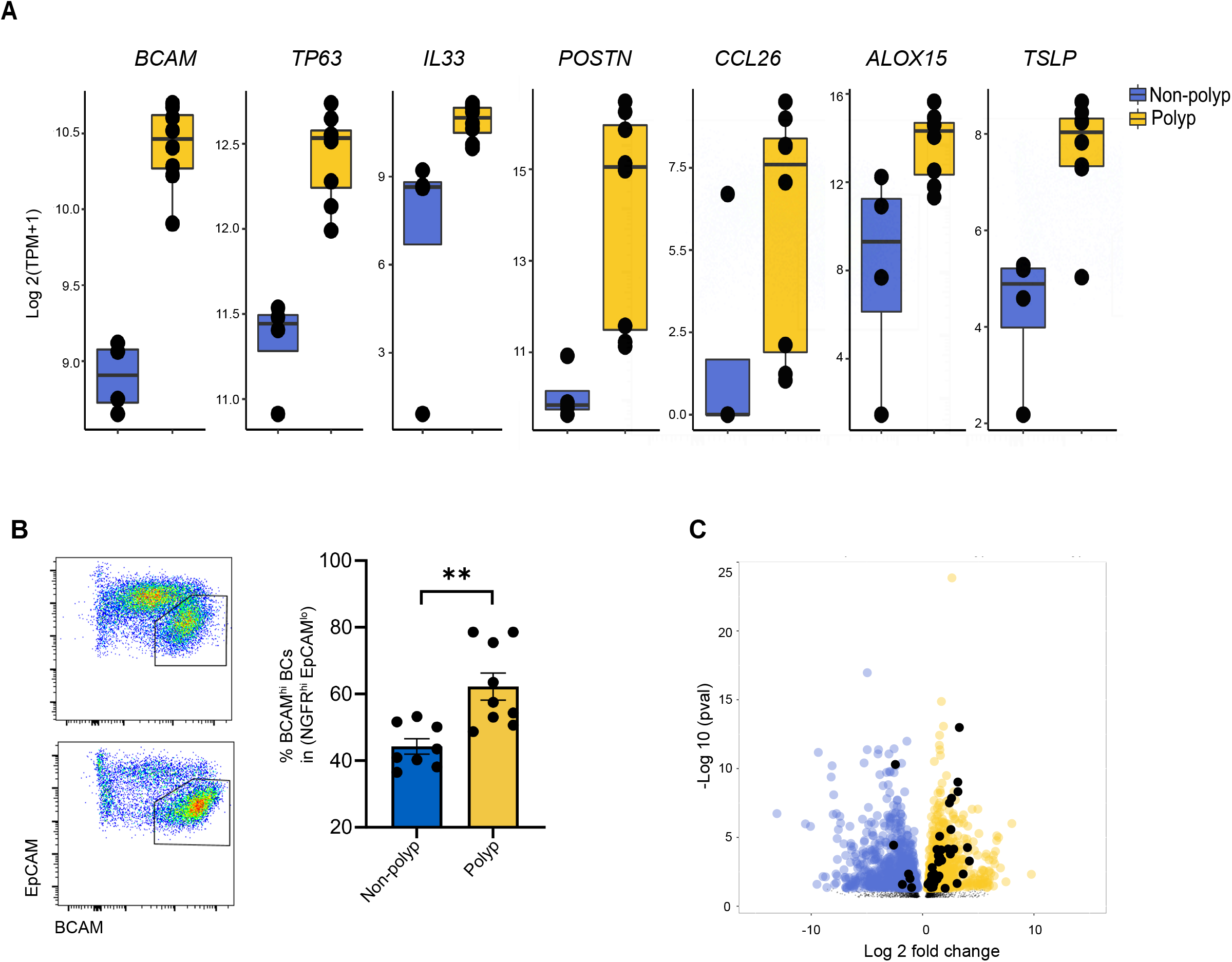
Expression of BCAM and the P63-dependent stem program is increased in CRSwNP and CRSsNP. (**A**) Expression of the indicated genes in bulk RNA-seq from human sinonasal polyp BCs and non-polyp BC controls. Adjusted p value (Benjamini-Hochberg) as follows: *BCAM* = 3.7e-13, *TP63* = 4.69e-7, *IL33* = 3.02e-2, *POSTN* = 5.20e-6, *CCL26* 3.44=e-1 (not significant), *ALOX15* = 2.95e-2, *TSLP* = 9.14e-07. (**B**) Flow cytometry on lin-EpCAM+ EpCs from CRSsNP and CRSwNP. Data are shown as mean ± SEM (** p<0.002; unpaired two-tailed t test). (**C**) Volcano plot showing differentially expressed genes up in CRSwNP (yellow) compared with CRSsNP (blue). The significant differentially expressed genes that are P63 target genes are colored in black.

To understand whether BCAM and TP63 were directly regulated by T2 cytokines, we cultured sinonasal BCs from healthy controls and from patients with CRSsNP or CRSwNP and treated them with either IL-4 and IL-13 or with TGF-β, which is known to induce EpC differentiation (59). In passaged unstimulated BC cultures, BCAM was expressed at high levels in all BCs with no BCAM^lo^ BC subset detected (**Fig. S5A**). Addition of TGF-β downregulated expression of BC markers NGFR and ITGA6, as expected, and also downregulated expression of *BCAM* (**Fig. S5B**). By contrast, IL-4/IL-13 upregulated BCAM expression but reduced NGFR and had little effect on ITGA6 (**Fig. 6A**). IL-4/IL-13 treatment also upregulated *TP63* and reduced expression of *HEY1*, indicating downregulation of Notch activity that is essential for BC differentiation into secretory EpCs (23, 31). Accordingly, we saw reduced *SCGB1A1* that marks secretory club cells (**Fig. 6B**). Additional markers of terminal EpC differentiation were not altered in this short-term experiment in submerged culture including *MUC5AC*, *FOXJ1*, *IL25*, and *POU2F3* (**Fig. S5C**). Taken together, these findings indicate that IL-4/13 reinforces a stem program in BCAM^hi^ BCs and *prevents* early steps in BC differentiation.

**Fig. 6.**
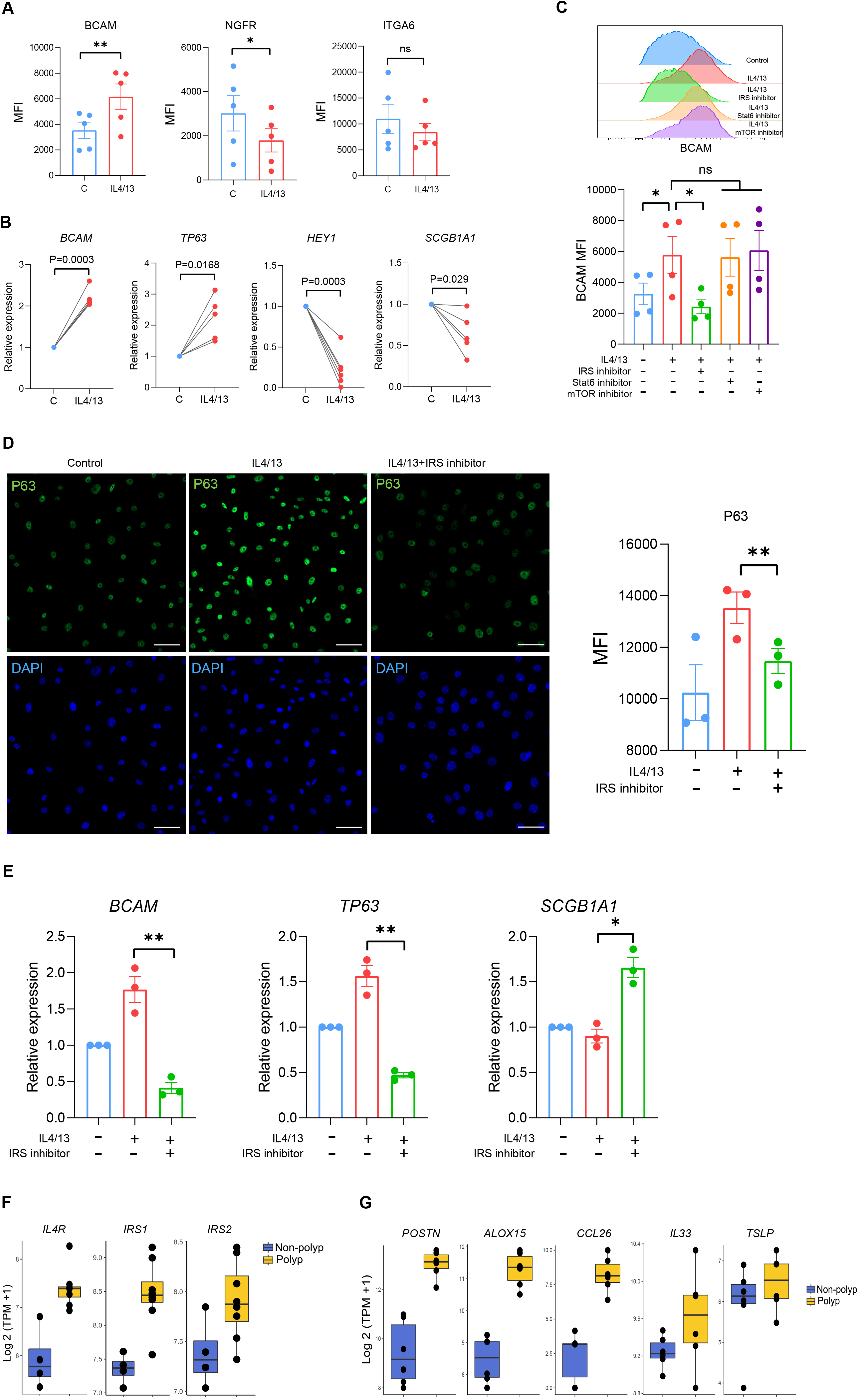
T2 cytokine- and IRS-dependent regulation of BCAM. (**A-E**) Human sinonasal BCs treated ± 10 ng/ml IL-4/13. (**A**) Mean fluorescence intensity (MFI) of BCAM, NGFR and ITGA6. Data are mean ± SEM (*p<0.05, ** p<0.01; paired two-tailed t test). (**B**) qPCR for the indicated genes. Data are mean ± SEM (*p<0.05, ** p<0.01; paired two-tailed t test). (**C**) Histogram (left) and MFI quantification (right) of BCAM expression in IL-4/13-treated BCs ± inhibitors of IRS-1/2, mTOR, and Stat6. Data are mean ± SEM (*p<0.05; paired two-tailed t test. (**D**) Representative images (left) and MFI quantification (right) of P63 staining on IL-4/13-treated BCs ± an IRS-1/2 inhibitor. Scale bar = 50 μm. Data are mean ± SEM (*p<0.05; paired two-tailed t test). (**E**) qPCR on IL-4/13-treated BCs ± an IRS-1/2 inhibitor. Data are mean ± SEM (*p<0.05, ** p<0.01; paired two-tailed t test). (**F**) Expression of components of IL-4R signaling pathway in bulk BC sequencing from CRSwNP or CRSsNP. *IL4R* and *IRS1* were significantly different (Benjamini-hochberg adjusted p-values < .05). (**G**) Expression of the indicated genes in scRNA-seq cluster 0 (BCAM^hi^ BCs) from CRSwNP and CRSsNP. Bonferonni threshold (1.59e-05) for *POSTN* = 3.21e-07, *CCL26* = 8.97e-06, and *ALOX15* = 1.28e-10.

IL-13 induces goblet cell metaplasia through a well-characterized STAT6-dependent pathway, but STAT6-independent pathways have previously been implicated in airway epithelial wound healing (60). Thus, we next assessed whether STAT6 or other IL-4Rα signaling pathways restrain BC differentiation. Pharmacologic inhibition demonstrated that IL-4/IL13-elicited BCAM expression was independent of STAT6 and mTOR signaling but dependent on IRS signaling (**Fig. 6C**). IRS inhibition also reduced P63 expression (**Fig. 6D**) and expression of BCAM and TP63 transcripts (**Fig. 6E**), while increasing expression of *SCGB1A1*. Assessment of transcripts for components of the IL-4R signaling pathway in bulk BC RNA-seq demonstrated that *IL4RA* and *IRS1* were upregulated in polyp BCs, as compared to controls (**Fig. 6F, Table S14**), while IRS2 showed a trend to increase that was not significant. Taken together, these data demonstrate that IL-4Rα and IRS signaling play an unexpected role in maintaining the BCAM^hi^ BC stem state, which accumulates in CRSwNP. Finally, we did interrogate our scRNA-seq dataset to assess whether BCAM^hi^ BCs from CRSwNP expressed higher levels of T2-inducible epithelial genes than BCAM^hi^ BCs from CRSsNP. We found that while *TSLP* and *IL33* were expressed at similar levels, *POSTN*, *CCL26*, and *ALOX15* were expressed more highly BCAM^hi^ BCs from CRSwNP than from CRSsNP.

## DISCUSSION

Identification of airway stem cells among plastic epithelial cell types is an important prelude to defining the molecular pathways that maintain stemness, promote normal tissue regeneration, and drive pathologic tissue remodeling. Previous studies have identified human and murine airway BCs as lin-EpCAM^lo^KRT5^+^NGFR^hi^ EpCs expressing ITGA6, PDPN, or GSIB4 (13, 18, 61–63) and detected significant heterogeneity within the BC compartment (4, 14, 24, 62, 64–68). However, cell surface markers to distinguish BC subsets and define BC biology have been lacking. Here we find that BCAM expression identifies molecularly and functionally distinct subsets of BCs in human and murine airways, with BCAM^hi^ BCs expressing high levels of P63 and exhibiting increased stem functions. Remarkably, BCAM^hi^ BCs are increased in the sinonasal mucosa of patients with the T2 inflammatory disease CRSwNP, and their P63 expression is upregulated through a T2 cytokine and IRS-dependent signaling pathway. These findings identify a robust marker of airway stem cells in mouse and human and define a T2 molecular pathway that promotes their persistence.

The top marker genes identified in BCAM^hi^ BCs include *BCAM*, the keratins *KRT5*, *KRT15*, *KRT17*; the canonical BC transcription factor *TP63* (19, 38) and its *S100A2* target (69–71); the stem cell regulator *MMP10* (72, 73); and diverse drivers of EpC proliferation including *LAMB3* (74), *MYC* (75), and the Yes-associated protein target *CYR61* (76). Each of these transcripts marks multipotent progenitor BCs from the lower airway (4, 14, 15, 68, 77). Moreover, analysis of differentially expressed genes between BCAM^hi^ and BCAM^lo^ BCs from the sinus and the lung (14) demonstrate that the distinct BCAM^hi^ and BCAM^lo^ gene programs seen in the sinonasal mucosa are largely retained in the bronchial tree. Additional support for this BCAM^hi^ vs BCAM^lo^ distinction in the tracheobronchial tree is found in our murine studies. Murine tracheal BCAM^hi^ BCs, but not BCAM^lo^ BCs, express high levels of Ki67 *in vivo*, demonstrate robust colony formation and wound healing in ex-vivo assays, and are rapidly labelled in lineage tracing studies. Additionally, we found that murine tracheal BCAM^hi^ BCs are enriched in *Wnt*, *Notch*, *Rho*, and *Trp63* pathway genes expected in airway stem cells. Taken together, these findings indicate that the BCAM^hi^ / BCAM^lo^ distinction detected in the sinonasal mucosa is likely to be useful in characterizing bronchial BCs.

BCAM is a member of the immunoglobulin superfamily, broadly expressed in erythroid, epithelial, endothelial, and smooth muscle cells. BCAM binds the alpha chain of the extracellular matrix protein laminin 5 (78, 79) to regulate cell adhesion and migration (80). Activation of BCAM promotes ERK/MAPK signaling, with an increase of RhoA and a decrease of Rac1 activity, that favors cell adhesion and colony formation in fibroblasts (81) and prevents biliary differentiation during liver regeneration (82). While RhoA signaling plays a central role in airway EpC differentiation (83, 84), whether BCAM is a critical regulator of BC RhoA functions will require further study. Additionally, BCAM binding to laminin α5 competitively inhibits integrin binding and alters integrin-mediated functions (85). This indicates an additional potential pathway by which BCAM expression can alter the regenerative functions of laminin α5 (86, 87) that are central to epithelial homeostasis and differentiation (88, 89).

Notably, we found that expression of *BCAM* and *TP63* were highly correlated in pseudotime, highly correlated across BCs from subjects with CRSwNP, and similarly induced by IL-4/13. Moreover, we found that human BCs from CRSwNP express higher levels of p63-dependent genes. CHIP-seq data demonstrate that p63 binds to a region in the BCAM promoter (90), and a recent study identified that overexpression of *Np63* in 293T cells increases luciferase activity in a reporter containing the BCAM promoter region (91). Thus, the reproducibility with which BCAM expression identifies airway stem cells across species and conditions may reflect its regulation by p63. Further studies are needed to understand this relationship, and to understand whether targeted inhibition of IL-4Rα reduces expression of p63 and BCAM *in vivo*. Previous studies have identified basal luminal progenitors (24, 31, 62), and KRT4/13+ hillock BCs (64, 66, 77) as descendants of KRT5+P63+ BCs that additionally express KRT8. In naïve murine trachea KRT8 was robustly expressed in the differentiated EpC gate, but it was not detected in either BCAM^hi^ or BCAM^lo^ BCs. Similarly, in human samples, we detected the highest levels of *KRT8* in transitional epithelial EpCs (clusters 2-4), which also expressed differentiation markers *HES1* and *SERPINB3*, but detected less in BCAM^hi^ and BCAM^lo^ BCs. Notably cluster 2 co-expressed *KRT5*, *TP63*, and *KRT8* and directly followed the BCAM^lo^ BC cluster in trajectory analysis. Furthermore, confocal analysis demonstrated that both BCAM^hi^ and BCAM^lo^ BCs were detected on the basement membrane, distinct from KRT8+ luminal BCs. Taken together, these data suggest that BCAM^hi^ and BCAM^lo^ BCs defined here represent earlier stages of differentiation than previously identified KRT8+ BCs.

Primary human sinonasal BCAM^lo^ BCs could not support the generation of air-liquid interface cultures *ex vivo* and murine BCAM^lo^ BCs had reduced *ex vivo* colony formation. This was unexpected, given numerous murine studies in the lower airway demonstrating the potential regenerative function of differentiated EpC subsets (25, 92–94) and our data in the murine trachea demonstrating inducible expression of Ki67, SSEA1, and Muc5AC in BCAM^lo^ BCs during injury. Taken together, our findings highlight the importance of local environmental cues in supporting replication that cannot be recapitulated *ex vivo*. As BCs passaged in submerged cultures from the upper and lower airway lose the distinction between BCAM^hi^ and BCAM^lo^ subsets, further studies using primary sorted human BCAM^lo^ BCs from the upper and lower airway in different culture systems will be needed to fully test the replicative potential of BCAM^lo^ BCs.

Here we found that the expansion of BCs in the T2 inflammatory disease CRSwNP that we previously reported (13) is due to an increase in the BCAM^hi^ BC subset. Moreover, we found that IL-4/13 upregulates the expression of BCAM and P63 in passaged BCs while reducing Notch activity and expression of *SCGB1A1*, which marks differentiated club EpCs. IL-4/13-elicited upregulation of BCAM and P63 was not mediated by STAT6 or mTOR signaling, but was reduced with an inhibitor of IRS-1 and IRS-2, each of which contributes to IL-4R-dependent functions in hematopoietic cells (95–97). While both IRS-1 and IRS-2 are required for IL-4-elicited migration of differentiated EpCs (60), only IRS-1 is significantly upregulated in polyp BCs. IRS-1 has a demonstrated role in promoting murine embryonic stem cell survival in vitro (98) and maintaining the Sox9+ intestinal stem cell pool *in vivo* (99), indicating that this is a likely mediator of IL-4/13-elicited stem function. Notably, a role for IL-4/13 in reinforcing stem function was not expected as prior studies have demonstrated that IL-4/13 drives the differentiation of EpCs to goblet cells (100, 101). Furthermore, T2 cytokine-driven maintenance of the BCAM^hi^ BC stem state has potential significant pathobiologic sequelae, as we find that many epithelial mediators of T2 inflammation including *IL33*, *TSLP*, *ALOX15*, and *CCL26* (102) are expressed in this cell type. Taken together, these data highlight the potential for T2 cytokines to drive a type of airway remodeling that in turn supports persistent disease.

This study identifies BCAM as a robust marker of airway stem cells using transcriptional, flow cytometric, and functional assays in murine trachea and human sinus tissue donated at surgery. Although we were able to compare our data to scRNA-seq data from the bronchial mucosa (14) to observe the global distinction between BCAM^hi^ and BCAM^lo^ BCs across tissues, functional analysis of primary bronchial BCAM^hi^ and BCAM^lo^ BCs will be needed to validate these observations, given the clear distinctions in the EpC compartment across the upper and lower airways (77). Additionally, although we find that high levels of BCAM distinguish multipotent progenitors among BCs, we do not have a similarly robust marker of murine BCAM^lo^ BCs. In the murine trachea, this group begins to express marks of secretory differentiated EpCs over the course of injury and repair, suggesting that additional markers are needed to distinguish this cell.

Together our data demonstrate 1) the transcriptional and functional distinctions between multipotent progenitor BCAM^hi^ BCs and their immediate BCAM^lo^ BC descendants, 2) the expansion of BCAM^hi^ BCs in a T2 high disease - CRSwNP, and 3) the IRS signaling pathway through which the T2 cytokines IL-4 and IL-13 can promote stem function. These findings highlight a role for T2 cytokines in the epithelium beyond canonical goblet cell metaplasia and support a growing literature demonstrating a role for T2 cytokines in tissue repair across diverse cell types (103–105). Moreover resolving the cell surface phenotype of these early airway precursors provides an opportunity to isolate and assess the molecular features of dysplastic BCs that are increasingly detected in a range of airway diseases from CRSwNP (13) to chronic obstructive pulmonary disease (106) and idiopathic pulmonary fibrosis (68, 107).

## AUTHOR CONTRIBUTIONS

Conceptualization: XW, NRH, JAB, MGA, NAB

Methodology: SR

Collection of human samples: AZM, RER, RWB, NB, KMB, DG, TR, and TML.

Human single cell and bulk sequencing analysis: NRH, XW, SS, SR, MGA, NAB

Murine bulk sequencing analysis: NRH, XW, MGA, NAB

Human *ex vivo* experiments and imaging: NRH, XW, ML

Murine *in vivo* and *ex vivo* experiments: XW, QY

Funding acquisition: JAB, NAB

Project administration: NAB

Supervision: MGA, NAB

Writing – original draft: XW, NRH, MAG, NAB

Writing – review & editing: JAB, TML, KMB, DFD, XW, NRH, MGA, NAB

## Abbreviations

T2: type 2
CRSwNP: chronic rhinosinusitis with nasal polyposis
CRSsNP: chronic rhinosinusitis sans nasal polyposis
EpCs: epithelial cells
BCs: basal epithelial cells
IL-4: interleukin-4
IL-13: interleukin-13
scRNA-seq: single cell RNA-sequencing
BCAM: basal cell adhesion molecule
KRT5: keratin 5
TP63: tumor protein 63
NGFR: nerve growth factor receptor
ITGA6: integrin alpha 6
EpCAM: epithelial cell adhesion molecule
GSIB4: Griffonia simplicifolia lectin B4
SCGB1A1: secretoglobin family 1A
SSEA1: stage-specific mouse embryonic antigen-1
muc5AC: mucin 5AC
STAT6: signal transducer and activator of transcription 6
IRS: insulin receptor substrate
mTOR: mammalian target of rapamycin

## ACKNOWLEGEMENTS

We wish to acknowledge Juying Lai for help with sample preparation, and Adam Chicoine for help with sorting.

## METHODS

### Study design

This study was designed to characterize basal cell subsets in the human and murine airway and to understand the alterations in BC programs in type 2 inflammatory diseases such as CRSwNP. This objective was addressed by: (i) reanalysis of scRNA-seq (13) of non-proliferating surface airway epithelial cells in surgical excisions of human sinonasal mucosa from patients with CRSwNP (n = 6) and CRSsNP (n = 6) to identify potential markers of BC subsets. This cohort was matched for age, sex, use of oral corticosteroids (none), and leukotriene-modifying agents at the time of surgery. (ii) flow cytometric analysis and *ex vivo* studies of primary human sinonasal BCs from CRS or healthy control subjects to demonstrate distinct BC functions; (iii) development of a murine flow cytometry panel, bulk RNA-seq datasets, BC lineage tracing system, and a model of airway damage and regeneration to characterize BC subsets in C57BL/6 mice; (iv) assessment of human BCAM^hi^ BCs *ex vivo* with and without type 2 cytokines. All in vitro and *in vivo* results are either representative of or pooled across three to five independent experiments.

### Single cell analysis of a public human sinonasal dataset

Seq-well data from 18546 cells (13) were initialized into a Seurat(citation) object after filtering out cells with less than 300 genes or over 12,000 UMIs. Gene UMI counts for each cell were normalized with a scale factor of 10,000 and natural log transformed with a pseudo count - log(counts +1). Normalized counts were centered and scaled for Principal Component Analysis (PCA), which was performed on the top 2000 genes ranked by their standardized variance using the Seurat function FindVariableFeatures. The top 20 Principal Components (PCs) were corrected with Harmony (17), integrating over donor and disease covariates. A neighborhood graph of cells (k-Nearest Neighbors(kNN)) was generated with the FindNeighbors function from Seurat (arguments). Cells were assigned to clusters using a resolution of 1.2 (Louvain algorithm) (**Fig. S1A-B**). Top marker genes for each cluster were generated using a Wilcoxon rank sum test and auROC with the Presto package (https://github.com/immunogenomics/presto) and clusters were annotated based on previous publication (13) (**Table S1**).

Sub-clustering of non-proliferating surface airway EpCs was performed in 3 subsequent iterations. Cells that were identified coarsely as Basal, Apical or Ciliated EpCs were subset into a new object, a new set of variable features was obtained, and the normalized counts re-scaled. PCA and Harmony corrections were re-run, again integrating for patient origin and disease status. Top 10 PCs were used for kNN classification and Louvain clustering with a resolution of 0.6. The first iteration resulted in an EpC sub-cluster (cluster 4) marked by high expression of immunoglobulin genes (**Fig. S1D far left panel**, **Table S2**). To determine whether or not underlying biology was driving this cluster, we removed immunoglobulin genes as possible drivers from the clustering analysis. In the second iteration, prior to re-scaling, immunoglobulin genes were removed from the variable feature matrix. The same parameters were used to cluster as in the first iteration. After harmony corrections and re-clustering, the previously identified cells did not cluster distinctly (**Fig. S1D, right and far right panels**). Thus, we considered these cells as contaminants with higher amounts of ambient transcripts from the digest. For the third iteration, cells expressing IG genes (cluster 4 from interation 1) were discarded. Subsequent steps to arrive at clustering were the same as the first iteration. 6970 cells were assigned to 9 clusters with good representation across donors and disease (**Fig. S1B-C**). Top markers were generated for each cluster with a Wilcoxon rank sum test and auROC statistic implemented in Presto.

### Trajectory Analysis

In order to define EpC differentiation, we performed trajectory analysis with the Slingshot package (108). A minimum spanning tree (MST) was constructed using Euclidean distances between cell clusters to build lineage topography and a principal curves algorithm generated smooth representations of each lineage. Each cell was assigned a pseudo time score via orthogonal projection onto each curve.

### Differential Gene Expression

The package glmGamPoi (109) was used to perform differential gene expression analysis. Briefly, a negative binomial generalized linear model was fit to each gene (genes were included in analysis if they were expressed in 10% of the smallest cluster). A quasi-likelihood ratio test as implemented in glmGamPoi was utilized to test for differential expression. For differential expression testing across pseudo time, scores generated by Slingshot for Lineage2 (secretory lineage), were rounded to the nearest whole integer and treated as a continuous variable. Sex, polyp status, and number of UMI’s were added as fixed effects to the model. For differential comparison between BCAM^hi^ and BCAM^lo^ BCs, a negative binomial generalized linear model was fit accounting for sex, polyp status and number of UMIs. BCAM^hi^ BCs were used as the reference level in the differential testing. For differential comparison between polyp vs non-polyp in BCAM^hi^ BCs in the sinonasal dataset, a negative binomial generalized linear model was fit accounting for sex, polyp status, and number of UMIs. Cells were aggregated across donor and tested as pseudo bulked samples. For all single cell comparisons, nominal p-values were adjusted with a Bonferroni correction to account for multiple-testing and results were considered significant when adjusted p-values were below 0.05.

### Single cell analysis of public bronchial dataset

A single cell dataset of healthy bronchial EpCs from brushings and biopsies (14) was downloaded and loaded into R. Original embeddings and cell annotations from paper were used for visualization and differential gene expression analysis. For differential comparison between bronchial BC1 and BC2 a negative binomial generalized linear model was fit accounting for source (tissue-site collection), number of UMIs, and donor.

### Mice

Animals treated with ALT were compared to untreated naïve mice. Age- and gender matched mice were randomly allocated to each experimental arm. No mice were excluded from study. Male and female mice ages 2 – 4 mos were used including WT C57BL/6 and ChAT^BAC^-eGFP [B6.Cg-Tg(RP23-268L19-EGFP)2Mik/J] mice. Krt5-^CreERT2^ mice (029155) and Ai9(RCL-tdT) (007909) were purchased from the Jackson Laboratory, Krt5^creERT2^: R26^tdTomato^ mice were generated by crossing Krt5-^CreERT2^ mice with Ai9(RCL-tdT) mice, adult heterozygous Krt5^creERT2^: R26^tdTomato^ mice were used in this study. 1 mg tamoxifen (T5648, Sigma) in sunflower seed oil (S5007, Sigma) was intraperitoneal injected in mice for 5 continuously day, intranasal sensitization was performed 7 days after the final tamoxifen injection.

### Intranasal sensitization and tissue preparation

After anesthesia with an intraperitoneal injection of ketamine (10 mg/kg) and xylazine (20 mg/kg), mice received intranasal application of 15 μg of *Alternaria* alternata culture filtrate antigen (362027, Greer Laboratories) in 20 μl of sterile PBS. This was repeated for the indicated number of weeks. Mice were euthanized at the indicated time points, trachea and lung tissues were harvested for histology or digested for flow cytometry. For preparing mouse nasal tissue immunofluorescent staining, after overnight fixation using 4% PFA, specimens were decalcified in 15% EDTA for 2 weeks at 4 C, after 30% sucrose overnight, specimens were imbedded in OCT for slide preparation.

### Mouse tracheal epithelial cell isolation for flow cytometry

Mouse tracheas were harvested, cleaned of connective tissue, opened longitudinally, and digested in prewarmed Tyrode's solution (PY-912, Boston BioProducts) containing Papain 8U/ml (P3125, Sigma), L-Cysteine 250 μg/mL (C7352, Sigma), Collagenase IV 700U/ml (LS004189, Worthington), and DNaseI 50 μg/mL (11284932001, Sigma) for 10 min at 37 ℃ with shaking at 200 rpm. Trituration with a 3 ml 18G syringe was performed 5 times, and digests were spun quickly (10-20 sec) to pellet undigested tissue. The supernatant was harvested into a new tube containing 25 μg/mL leupeptin (L2023, Sigma) and 2 μM EDTA (E6758, Sigma) to stop the reaction, mixed well, and placed on ice. Undigested tissue fragments were incubated with fresh digestion buffer as above 1-2 more times. All cell suspensions were combined and filtered through 100 um strainers (352360, Fisher Scientific). The filtered digests were centrifuged at 1000g for 10 min, and cell pellets were resuspended with cold FACS buffer for downstream analysis.

### Flow cytometry

For mouse tracheal BCs, single cell suspensions were blocked with Fc-blocker (101320, Biolegend) for 10 min on ice, then incubated with appropriate antibodies: BCAM (D295-3, MBL International), EpCAM (118213, Biolegend), CD31 (102434, Biolegend), CD90 (105328, Biolegend), CD45 (103116, Biolegend), GSIB4 (L2895, Sigma), CD24 (101821, Biolegend), SSEA1 (125613, Biolegend), NGFR (ab52987, ABCAM) and CD49f (313623, Biolegend). 7AAD (420403, Biolegend) was added immediately before cell aquisition. For intracellular staining of SCGB1A1 (ab40873, Abcam), KRT5 (ab224984, ABCAM) and KRT8 (ab192468, Abcam), cells were fixed with fixation buffer (420801, Biolegend), then permeabilized with permeabilization wash buffer (421002, Biolegend) following the manufacturer’s protocol. For nuclear staining of P63 (ab124762, Abcam) and IL-33 (AF3626, R&D), the True-Nuclear™ transcription factor buffer set (424401, Biolegend) was used following the manufacturer’s protocol. After excluding debris and dead cells, EpCAM^+^ Lineage-(CD31, CD90 and CD45) EpCs were gated into 3 population with BCAM and GSIB4: BCAM^hi^ BCs (BCAM^hi^ GSIB4^+^), BCAM^lo^ BCs (BCAM^lo^ GSIB4^+^) and Diff EpCs (BCAM^lo^ GSIB4^−^). For human nasal EpCs, cells were stained as above using EpCAM (324219, Biolegend), CD45 (304035, Biolegend), CD31(303113, Biolegend), CD90 (3228125, Biolegend), BCAM (566661, BD), NGFR (345109, Biolegend), ITGA6 (313628, Biolegend). Data were acquired on BD FACSCanto II or LSRFortessa flow cytometer, flow cytometric analysis was performed using FlowJo.

### qPCR

Briefy, RNA was collected using RNeasy Mini Kit (74004, Qiagen) according to the manufacturer’s instruction. After synthesis of cDNA using SuperScript III First-Strand (18080-400, Invitrogen) under T100 Thermal Cycler (Bio-RAD, qPCR was performed using the Mx3000/Mx3005P Real-Time PCR System. All primers were used in this study were ordered from Qiagen BCAM (PPH12956A), SCGB1A1 (PPH01032F), P63 (PPH00177F).

### Histological assessment

Trachea samples were harvested and fixed with 4% PFA overnight, the tissue was wash with PBS for 3 time (10 min each), followed by paraffin embedding and sectioned with 6 μm thickness. H&E staining were performed using standard protocol.

### Immunofluorescence staining

Mouse trachea and human ALI culture samples were harvested and fixed with 4% PFA (BM-155, Boston BioProducts) overnight at 4 °C, the tissue was rinsed with PBS for 3 time (10 min each), incubated in 30% sucrose overnight at 4 °C. Mouse trachea and human ALI cultures were then imbedded with Tissue-Plus™ O.C.T. Compound (23-730-571, Fisher Healthcare) and cryosectioned at 5 μm thickness using a cryostat (Leica CM1850) and adhered to positively charged glass slides (12-550-15, Fisher Scientific). Cryosections were permeabilized with 0.2% Triton X-100 for 20 min and washed with PBS for 10 min, then blocked in 1× blocking buffer (ab126587, ABCAM) for 1h at room temperature. Afterwards, tissue samples were incubated with appropriate primary antibodies overnight at 4℃, washed 3 times with PBS, incubated with secondary antibodies for 1h at room temperature, washed 3 times with PBS, and mounted with mounting medium containing DAPI (ab104139, ABCAM), samples are ready for visualization under a microscope. Primary antibodies used on mouse experiments were: BCAM (AF8299, R&D), Muc5ac (ab3649, Abcam), Acetylated tubulin (T6793, Sigma), SCGB1A1 (ab40873, Abcam), Ki67 (14-5698-82, Invitrogen), KRT5 (ab52635, Abcam), P63 (GTX102425, Gene Tex). Primary antibodies used on human experiments were: BCAM (566661, BD), KRT5 (ab52635, Abcam), P63 (ab124762, Abcam), Muc5ac (ab3649, Abcam), IL33 (AF3625, R&D), KRT8 (ab53280, Abcam), Acetylated tubulin (T6793, Sigma). Secondary antibodies were used: Alexa Fluor conjugates (488, 594 and 647) were used at 1:500 dilution (Invitrogen). Images were obtained with a Zeiss LSM 700 Laser Scanning Confocal Microscopy, images were analyzed and merged using ImageJ (National Institutes of Health, Bethesda, MD).

### Mouse tracheal EpCs expansion

Submerged culture. Cell culture plates were precoated with 0.03 mg/ml rat tail collagen (354236, Corning) for 2h. Tracheal epithelial cells isolated from mice were seeded into the precoated plate with Keratinocyte SFM medium (17005042, Thermo Fisher Scientific) containing 1% P/S, 0.1% amphotericin B (15290, Thermo Fisher Scientific), 25 ng/ml EGF (E4127, Sigma), 30ug/ml BPE (13028-014, Thermo Fisher Scientific), 1μM Isoproterenol (16504, Sigma), 10 μM Y-27632 (10005583, Cayman) and 5 μM DAPT (13197, Cayman). After several days of culture when BCs were 90% confluent, they were lifted with Accutase (A11105, Gibco) and passaged or saved in liquid nitrogen. Stored BCs were seeded into 24 well plates. After 70% confluency was achieved over days of culture, the medium was changed. Flow cytometry on BCAM, NGFR and ITGA6 was measured after 48 hrs culture in various conditions.

### Mouse tracheal BCAM^hi^ and BCAM^lo^ BC sorting and colony forming assay

Mouse tracheas were harvested, cleared of connective tissue, opened longitudinally and digested overnight at 4℃ in Opti-MEM (31985-070, Gibco) containing 1% P/S, 0.1% amphotericin B and 0.15% pronase (10165921001, Roche Diagnostic). The single cell suspension was harvested and stained to identify BCAM^hi^ and BCAM^lo^ BCs, and BCs were sorted into tdTomato^+^ GSIB4^+^ BCAM^hi^ BCs and tdTomato^+^ GSIB4^+^ BCAM^lo^ BCs using 5-laser BD FACS Aria Fusion cell sorter. Sorted BCAM^hi^ and BCAM^lo^ BCs were seeded directly into 6 well plate (2,500 cells/well) for submerged culture. After 5 to 7 days, the plate was fixed, and wells were scanned for tdTomato signal using GE Healthcare IN Cell Analyzer 2200. After scanning, the plate was stained for p63 and ki67, and images were acquired on a Zeiss LSM 700 Laser Scanning Confocal Microscopy.

### Wound healing assay on BCAM^hi^ and BCAM^lo^ BCs

Sorted BCAM^hi^ and BCAM^lo^ BCs were seeded into the Culture-Insert 4 well plate (80466, IBIDI) with submerged culture medium. After 90% confluency, the culture insert was removed, a scratch was generated, and images of the cell-free gap were acquired at the indicated times. The percentage of wound closure was calculated using ImageJ software.

### Human nasal EpC submerged culture

Sinus tissue was collected at the time of elective endoscopic sinus surgery from patients with CRSwNP and CRSsNP with the diagnosis based on established guidelines (110). The CRSwNP group included some patients with aspirin-exacerbated respiratory disease but no distinctions were made between these subtypes. Non-CRS control patients were undergoing sinus surgery to correct anatomic abnormalities by removal of concha bullosa. Origin of BCs for each assay is found in **Table S3**. Sinonasal tissue resections were chopped, digested in RPMI-1640 with 10% FBS containing Type IV collagenase (LS004189, Worthington) and DNaseI (10104159001, Sigma) with magnetic stir bar at 600 RPM at 37℃ for 30 min, triturated using a 25 ml syringe with a 16g needle every 15 min. The resultant cell suspension was filtered over a 70 um cell strainer, spun down at 500g for 10 min, and washed. 75 mm tissue culture flasks were precoated with 0.03 mg/ml collagen (4902, STEMCELL) for 2 h. Single cell suspensions were seeded into precoated 75mm flask with PneumaCult Ex basal medium: PneumaCul-Ex Medium (05008, STEMCELL) containing 500uL StemCell Hydrocortisone (07925, STEMCELL), 1% Penicillin-Streptomycin (15140122, Thermo Fisher Scientific), 0.1% Gentamycin/AmphotericinB, 1 μM A-83-01 (2939, Tocris Bioscience), 0.2 μM DMH-1 (4126, Tocris Bioscience), 5 μM Y27632, and 2.5 μM DAPT. After cell expansion, BCs were seeded into a 24 well plate. 24 hrs after seeding, the supernatant was removed and the cells were treated with and without 10 ng/ml IL-4/13 or 10 ng/ml TGFβ for 48 h. BCs were then collected for flow cytometry. For inhibition of IRS, mTOR and Stat6 signaling, cells were first treated with an IRS-1/2 inhibitor (S8228, 5um, Selleck Chemicals), an mTOR inhibitor (S2827, 50nm, Selleck Chemicals), or a Stat6 inhibitor (S8685, 50nm, Selleck Chemicals) for 1 h before addition of IL-4/IL-13. 48 h later, cells were collected for flow cytometry, qPCR, and imaging.

### Human BCAM^hi^ and BCAM^lo^ BC sorting and ALI cultures

Single cell suspensions were stained with CD31, CD90, CD45, NGFR and BCAM. After excluding debris and dead cells, lin-EpCAM^lo^NGFR+ EpCs were sorted into BCAM^hi^ BCs and BCAM^lo^ BCs. The sorted cells were seeded into a 6-well plate under submerged culture conditions. After several days of culture and reaching 90% confluency, BCs were detached and seeded in 6.5 mm Transwell plates (3470, Costar) at a density of 100k cells per well with PneumaCult Ex basal medium. The media in the upper and lower chambers were changed daily. After confluence was achieved, the media in the upper chamber was aspirated and ALI medium was added to the lower chamber every other day. ALI medium components were as follows: ALI Complete Base Medium (05001, STEMCELL Technologies) containing 100uL of PneumaCult™-Ex Plus 50X Supplement, 50uL StemCell Hydrocortisone (07925, STEMCELL Technologies), and 20 uL StemCell Heparin (07980, STEMCELL Technologies). After 21 days, the membrane insert was collected, washed with PBS, and fixed with 4% PFA for 15 min.

### Low-input RNA-seq of murine epithelial cells

Mouse trachea epithelial cells were sorted by 5-laser BD FACS Aria Fusion cell sorter, detailed gating strategy could be found in Fig. S2. In brief, after removing debris, doublets, and dead cells, EpCAM^+^ Lineage^−^ EpCs were sorted into FACS tube, then re-sorted the sample into three populations: BCAM^hi^ GSIB4^+^ (BCAM^hi^ BCs), BCAM^lo^ GSIB4^+^ (BCAM^lo^ BCs) and BCAM^−^ GSIB4^−^ (Diff). 1000 cells were collected for each population, mixed with 5ul TCL buffer (1031576, Qiagen) buffer directly and stored at −80C. Libraries were prepared using Smart-seq2 and 38bp paired-end sequencing was performed by the Genomics Platform of Broad Institute of MIT and Harvard.

Reads were pseudo-aligned and transcript expression quantified with Kallisto (citation) using GRCm39 (GCA_000001635.9). Quantification files were processed to raw counts with the tximport-DESeq2 pipeline. Samples were considered high quality for downstream analysis if greater than 12,000 genes and more than 10^6 raw counts were detected and if the samples had >99% of common genes expressed. PCA confirmed samples separated by sorted subsets, and there were no outliers. Differential gene expression analysis was performed with the package glmGamPoi, where a negative-binomial generalized linear model was fit with BCAM status and batch as fixed effects. A quasi-likelihood test implemented in glmGamPoi was used for differential testing across the BCAM status variate. Adjusted p-values to control for multiple testing correction with Benjamini-Hochberg were considered significant below .05.

### Analysis of bulk sequencing from human BCs

The abundance matrix (13) was downloaded (RSEM). Abundance estimates were rounded to whole integers and treated as raw counts to use for differential gene expression analysis. Differential gene expression analysis was performed with the package glmGamPoi, where a negative-binomial generalized linear model was fit with disease as the variable of interest. A quasi-likelihood test implemented in glmGamPoi was used for differential testing across the BCAM status variate. Adjusted p-values to control for multiple testing correction with Benjamini-Hochberg were considered significant below 0.05. For correlation analysis with BCAM, Pearson correlation was performed between pairs of genes from samples collected from patients with Nasal Polyps.

## Data and code availability

Low-input RNA-seq data has been deposited in GEO, GSE197274. CRSwNP single cell dataset has been deposited in ImmPort SDY1877. The public dataset from healthy lung was downloaded from https://www.covid19cellatlas.org/index.healthy.html#vieira19-bronchi (14). Code for main analyses will be available at https://github.com/nils-hallen/Single-Cell-Analyses.

## Statistics

Analysis was performed with GraphPad Prism software (version 9.3.1, GraphPad, La Jolla, CA) and R (version 4.03). Data indicate means ± SEMs in all bar graphs. P < 0.05 was considered significant. Other statistical analyses as detailed in Methods sections above and in Fig. legends.

## Study Approval

The Mass General Brigham Institutional Review Board approved the study and all subjects provided written informed consent prior to participation. The use of mice for this study was in accordance with review and approval by the Animal Care and Use Committee of Brigham and Women’s Hospital.

