## Supplementary material for "Type 2 Inflammation Drives an Airway Basal Stem Cell Program Through Insulin Receptor Substrate Signaling": Suppl Materials

**Supplementary Materials**

1. Supplementary Figures and Legends.
   a. Fig. S1: Bioinformatic and flow cytometric approach to characterize human BC diversity.
   b. Fig. S2: Assessment of $BCAM^{hi}$ and $BCAM^{lo}$ BCs in naïve murine airway.
   c. Fig. S3: Flow cytometric and confocal analysis of EpCs in ALT model of tracheal injury and repair.
   d. Fig. S4: Isolation of purified $BCAM^{hi}$ and $BCAM^{lo}$ BCs and gene expression profiles.
   e. Fig. S5: Analysis of BC surface markers in passaged and stimulated BCs.
2. Excel file with data Tables E1-14.

| | |
|---|---|
| Table S1: | Top markers of parent clustering with harmony |
| Table S2: | Top markers of epithelial clusters prior to removal of the IgJ+ cluster |
| Table S3: | Origin of BCs used for each assay |
| Table S4: | Top markers of final epithelial re-clustering |
| Table S5: | Significant genes that increase with pseudotime |
| Table S6: | Significant genes that decrease with pseudotime |
| Table S7: | Differentially expressed genes between BCAM Hi vs BCAM Lo |
| Table S8: | Genes shared between BCs in bronchial and sinonasal epithelium |
| Table S9: | Top 50 genes driving PC1 in the positive and negative direction |
| Table S10: | Genes enriched in murine BCAM Hi BCs |
| Table S 11: | Genes enriched in murine BCAM Lo BCs |
| Table S12: | Transcription factors differentially expressed across murine BC subsets |
| Table S13: | Pearson correlation of all genes with BCAM in sorted sinonasal BCs |
| Table S14: | Genes differentially expressed in BCAM Hi BCs from Polyp vs Non-polyp |

### SUPPLEMENTARY FIGURES

#### Fig. S1

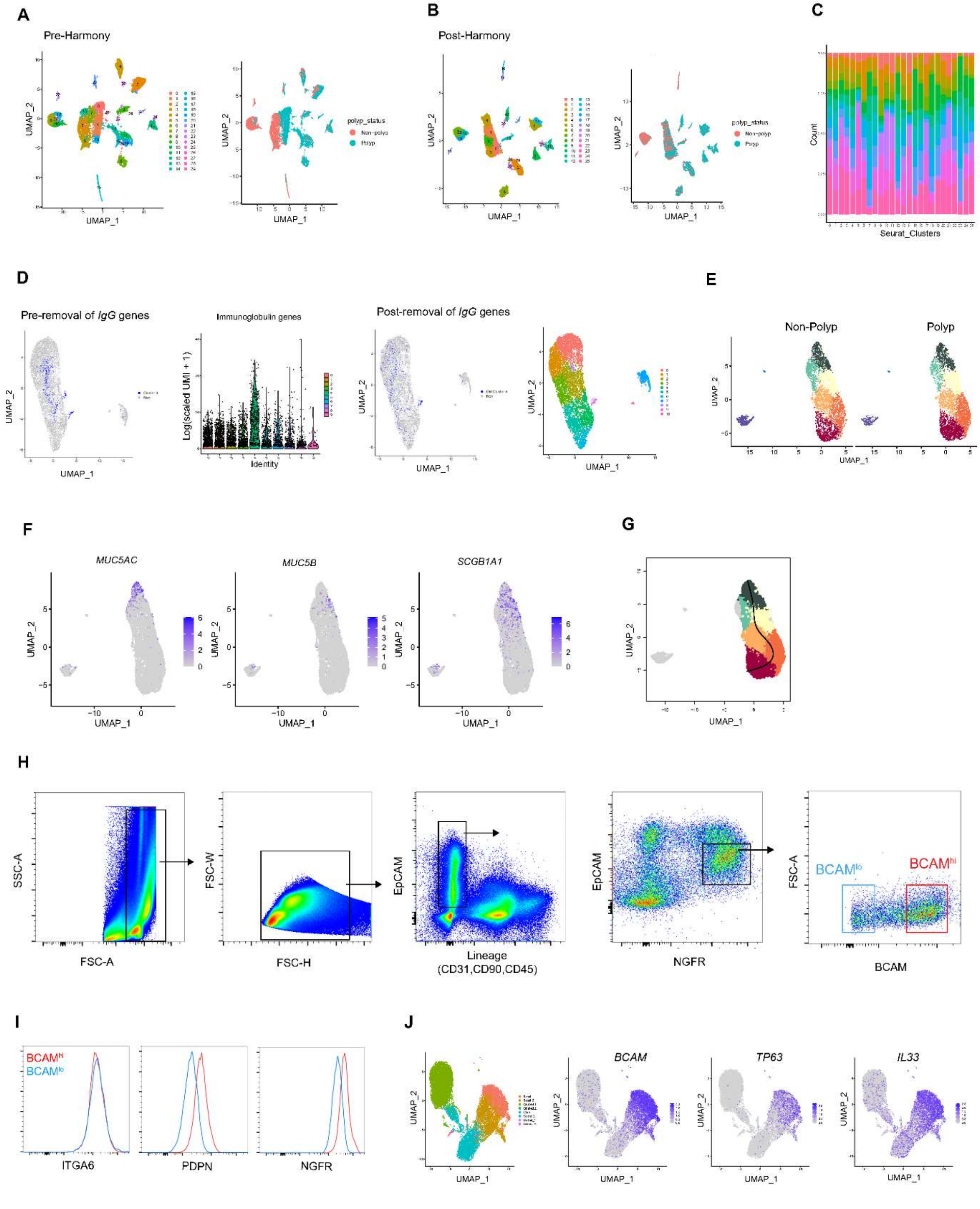

**Fig. S1: Bioinformatic and flow cytometric approach to characterize human BC diversity.** (**A**) UMAP of sinonasal surgical resections (18546 cells) from CRSwNP and CRSsNP (*13*) colored either by Louvain clustering (resolution, left panel) or disease status (right panel) prior to integration with Harmony (*17*). (**B**) UMAP of sinonasal surgical resections (18546 cells) from CRSwNP and CRSsNP colored either by Louvain clustering (resolution, left panel) or disease status (right panel) after integration with Harmony. (**C**) Distribution of cells from each donor across all clusters after integration with Harmony. (**D**) Analysis of reclustered surface airway epithelial cells (excluding proliferating BCs and submucosal gland cells). Feature plot of cluster 4 (far left panel) marked by high expression of immunoglobulin genes (iteration 1: Methods). Violin Plot of summed normalized counts of immunoglobulin genes across clusters (left panel). Feature plot demonstrating dispersion of cells from cluster 4 after removal of immunoglobulin genes from the variable feature matrix (iteration 2: Methods) (right panel). UMAP colored by Louvain clustering (resolution, iteration 2: Methods) (far right panel) after removal of cells expressing immunoglobulin genes. (**E**) UMAP visualization split by disease status. (**F**) UMAP colored by expression of key marker genes for secretory EpC subtypes. (**G**) UMAP colored by all cells assigned to the secretory lineage (lineage 2). The principal curve for lineage 2 is overlaid. (**H**) Gating strategy for flow cytometric sorting of BCAM$^{hi}$ and BCAM$^{lo}$ BCs from human sinonasal samples. (**I**) Flow cytometric evaluation of BC markers in BCAM$^{hi}$ and BCAM$^{lo}$ BCs demonstrating that both subsets express high levels of ITGA6, PDPN, and NGFR. (**J**) UMAP of bronchial EpCs from healthy controls (*14*) colored by pre-annotated cell identities (far left panel) and the indicated genes (left, right, and far right panels).

Fig.S2

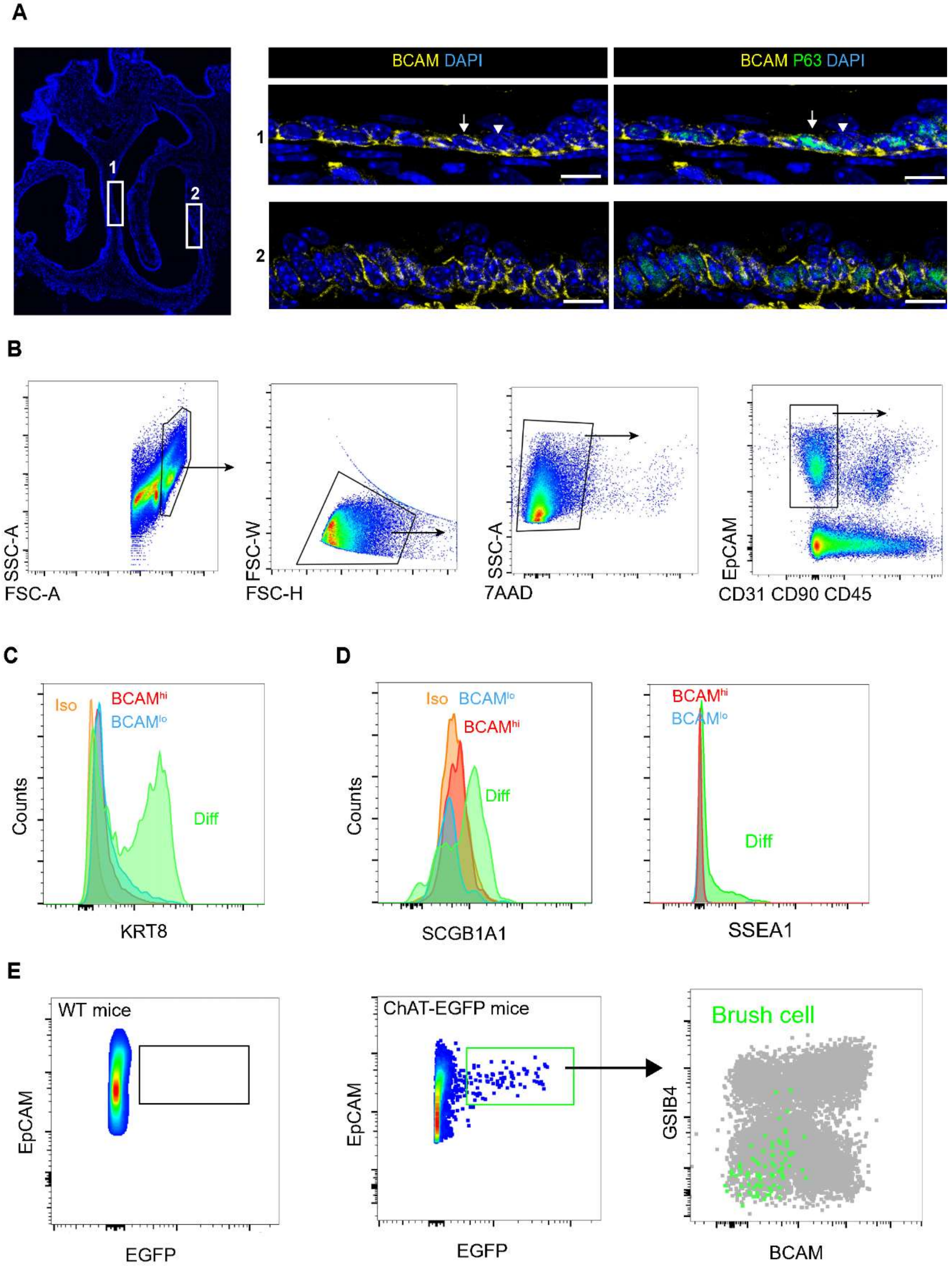

A
BCAM DAPI
BCAM P63 DAPI
1
2
B
SSC-A
FSC-A
FSC-W
FSC-H
SSC-A
7AAD
EpCAM
CD31 CD90 CD45
C
Iso
BCAM^hi
BCAM^lo
Diff
Counts
KRT8
D
Iso
BCAM^lo
BCAM^hi
Diff
Counts
SCGB1A1
BCAM^hi
BCAM^lo
Diff
SSEA1
E
WT mice
EpCAM
EGFP
ChAT-EGFP mice
EpCAM
EGFP
Brush cell
GSIB4
BCAM

**Fig. S2: Assessment of BCAM$^{hi}$ and BCAM$^{lo}$ BCs in naïve murine sinonasal mucosa and trachea.** (**A**) Representative immunostaining in naïve murine nasal tissue from the septum (1) and sinus (2). The scale bar represents 50 μm. (**B**) Flow cytometric gating to identify EpCs in the naïve murine trachea. After EpCAM$^{+}$CD31$^{-}$CD90$^{-}$CD45$^{-}$ EpCs were defined, cells were further gated into BCAM$^{hi}$ GSIB4$^{+}$ BCs (BCAM$^{hi}$ BCs), BCAM$^{lo}$ GSIB4$^{+}$ BCs (BCAM$^{lo}$ BCs), and BCAM$^{lo}$ GSIB4$^{-}$ differentiated EpCs (Diff) (**Fig. 2A**). (**C-D**) Flow cytometric staining for KRT8 (C) and SCGB1A1 and SSEA1 (D) expression in BCAM$^{hi}$ BCs, BCAM$^{lo}$ BCs and Diff EpCs. (**E**) Flow cytometric staining for EGFP in WT (left) and Chat-EGFP mice (middle panel) demonstrating rare EGFP$^{+}$ brush/tuft cells that fall in the GSIB4$^{-}$ Diff gate (right panel).

Fig. S3

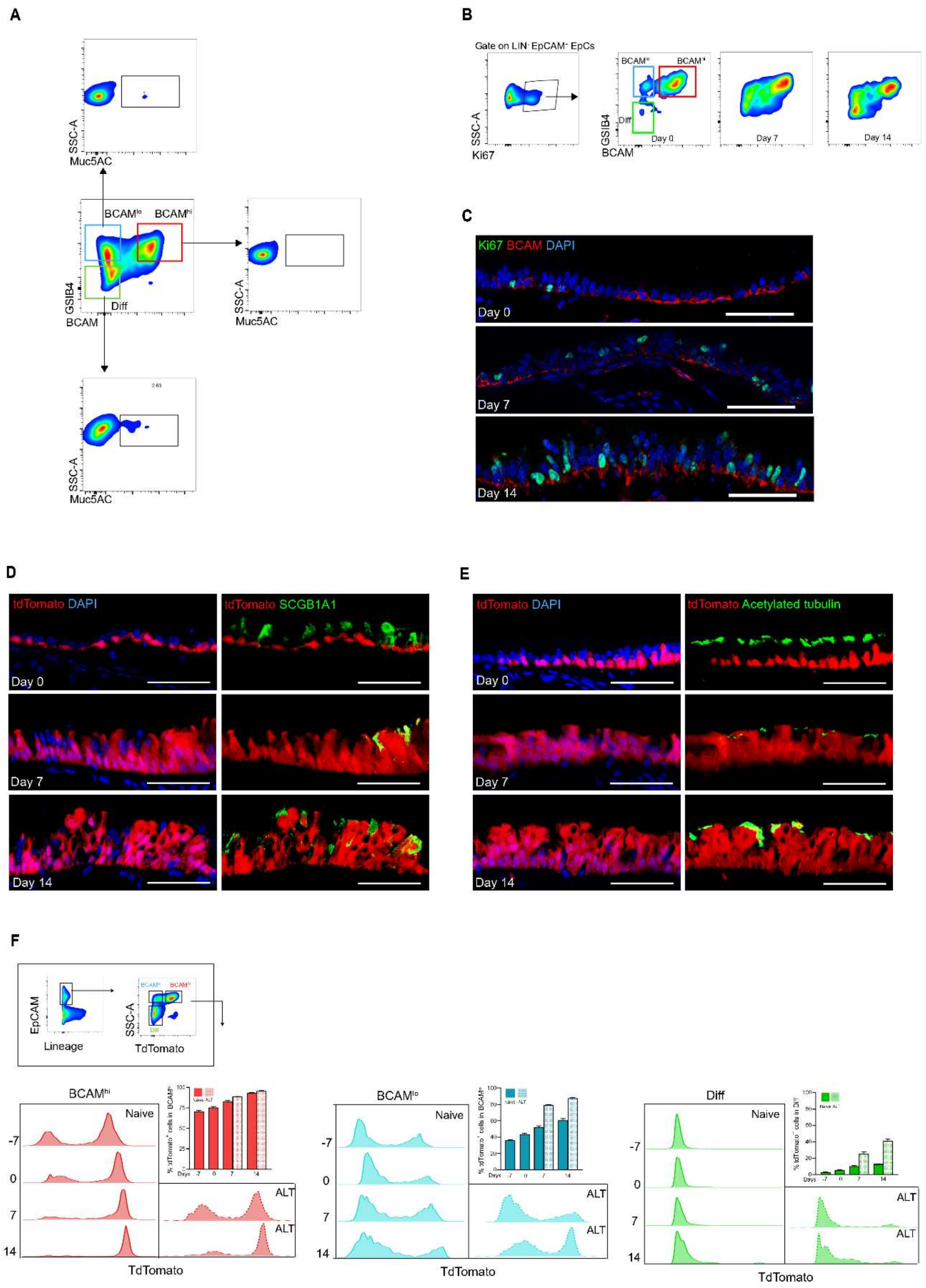
A
SSC-A
Muc5AC
BCAM^lo
BCAM^hi
GSIB4
BCAM
Diff
B
Gate on LIN- EpCAM+ EpCs
SSC-A
Ki67
GSIB4
BCAM
Day 0
Day 7
Day 14
C
Ki67 BCAM DAPI
Day 0
Day 7
Day 14
D
tdTomato DAPI
tdTomato SCGB1A1
Day 0
Day 7
Day 14
E
tdTomato DAPI
tdTomato Acetylated tubulin
Day 0
Day 7
Day 14
F
EpCAM
Lineage
SSC-A
TdTomato
Diff
BCAM^hi
BCAM^lo
Diff
Naive
ALT
-7
0
7
14
TdTomato
Days

**Fig. S3: Flow cytometric and confocal analysis of EpCs in ALT model of tracheal injury and repair.** (**A**) Flow cytometry gating and stain for Muc5AC. (**B**) Flow cytometry gating method for Ki67+ EpCs. (**C**) Immunofluorescence staining for Ki67 (green) and BCAM (red) in mouse trachea on day 0, 7 and 14. The scale bars represent 50 μm. (**D-E**) Lineage tracing of $KRT5^{+}$ BCs on day 0, 7 and 14. Trachea sections were co-stained with SCGB1A1 (D) and Acetylated Tubulin (E). The scale bars represent 50 μm. (**F**) Representative staining and percent of $BCAM^{hi}$ BCs (left), $BCAM^{lo}$ BCs (middle) and differentiated EpCs (right) that are labelled by TdTom in naïve and ALT at the indicated timepoints. Data are shown as mean ± SEM (n = 3).

**Fig. S4**

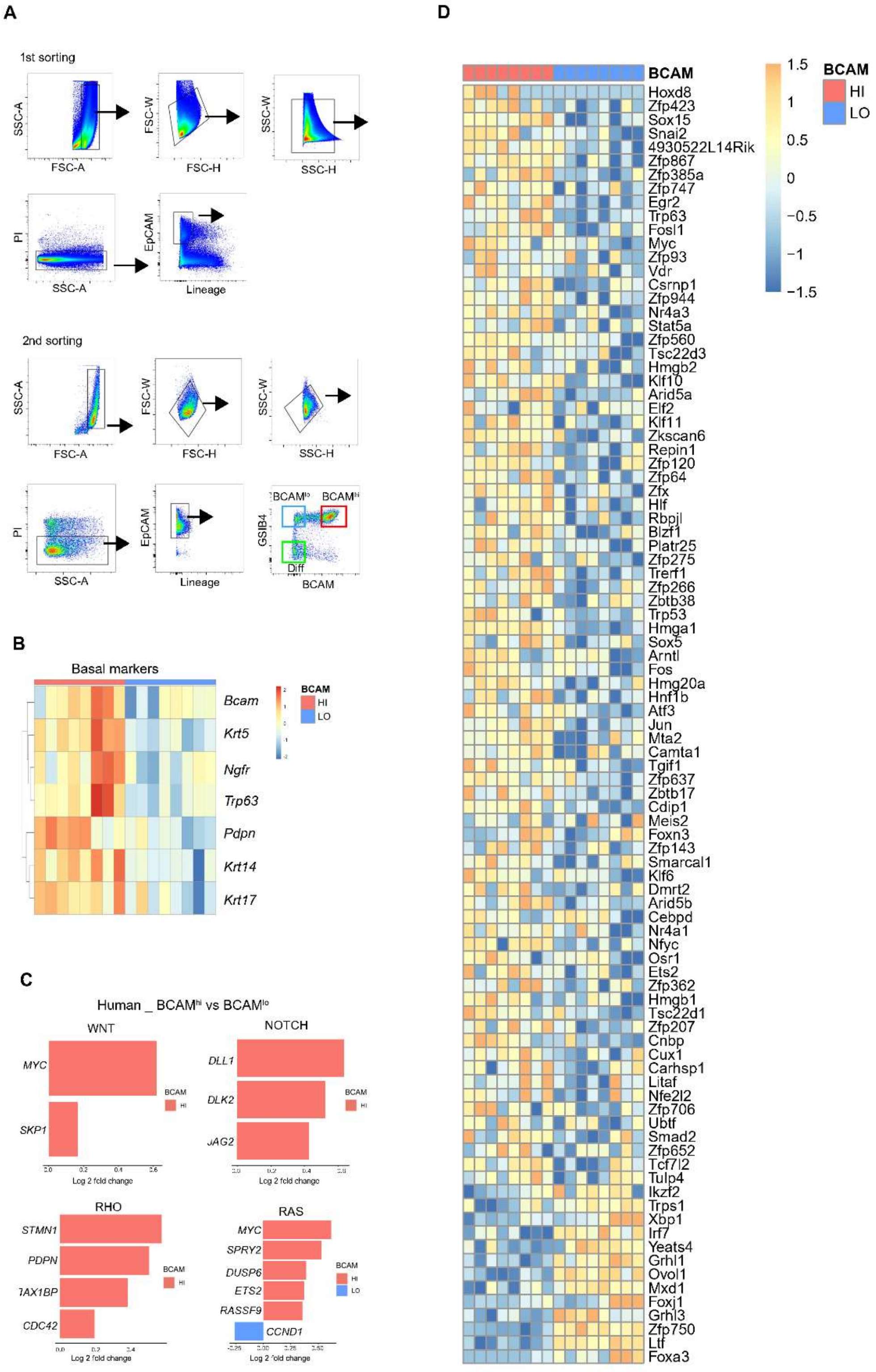
A
1st sorting
SSC-A
FSC-A
FSC-W
FSC-H
SSC-W
SSC-H
PI
SSC-A
EpCAM
Lineage
2nd sorting
SSC-A
FSC-A
FSC-W
FSC-H
SSC-W
SSC-H
PI
SSC-A
EpCAM
Lineage
BCAMlo
BCAMhi
GSIB4
Diff
BCAM
B
Basal markers
Bcam
Krt5
Ngfr
Trp63
Pdpn
Krt14
Krt17
BCAM
HI
LO
C
Human _ BCAMhi vs BCAMlo
WNT
MYC
SKP1
NOTCH
DLL1
DLK2
JAG2
RHO
STMN1
PDPN
CDC42
RAS
MYC
SPRY2
DUSP6
ETS2
RASSF9
CCND1
Log 2 fold change
BCAM
HI
LO
D
BCAM
1.5
1
0.5
0
−0.5
−1
−1.5
HI
LO
Hoxd8
Zfp423
Sox15
Snai2
4930522L14Rik
Zfp867
Zfp385a
Zfp747
Egr2
Trp63
Fosl1
Myc
Zfp93
Vdr
Csrnp1
Zfp944
Nr4a3
Stat5a
Zfp560
Tsc22d3
Hmgb2
Klf10
Arid5a
Elf2
Klf11
Zkscan6
Repin1
Zfp120
Zfp64
Zfx
Hlf
Rbpjl
Blzf1
Platr25
Zfp275
Trerf1
Zfp266
Zbtb38
Trp53
Hmga1
Sox5
Arntl
Fos
Hmg20a
Hnf1b
Atf3
Jun
Mta2
Camta1
Tgif1
Zfp637
Zbtb17
Cdip1
Meis2
Foxn3
Zfp143
Smarcal1
Klf6
Dmrt2
Arid5b
Cebpd
Nr4a1
Nfyc
Osr1
Ets2
Zfp362
Hmgb1
Tsc22d1
Zfp207
Cnbp
Cux1
Carhsp1
Litaf
Nfe2l2
Zfp706
Ubtf
Smad2
Zfp652
Tcf7l2
Tulp4
Ikzf2
Trps1
Xbp1
Irf7
Yeats4
Grhl1
Ovol1
Mxd1
Foxj1
Grhl3
Zfp750
Ltf
Foxa3

**Fig. S4: Isolation of purified BCAM$^{hi}$ and BCAM$^{lo}$ BCs and gene expression profiles.**
(**A**) Flow cytometric staining of murine EpC subsets for sorting and sequencing. (**B**) Heatmap of selected canonical BC marker genes that were significantly differentially expressed (Benjamini-hochberg adjusted p-values < .05) between murine BCAM$^{hi}$ and BCAM$^{lo}$ BCs. (**C**) Significant differentially expressed genes (Benjamini-Hochberg p-value < .05) from Wnt, Notch, Rho/Rock, and Ras signaling pathways in the human scRNA-seq data. These genes, first identified in BCAM$^{hi}$ murine BCs, are increased in BCAM$^{hi}$ human BCs with the exception of *CCND1*. (**D**) Heatmap of transcriptional factors differentially expressed (Benjamini-hochberg adjusted p-values < .05) between BCAM$^{hi}$ and BCAM$^{lo}$ murine BCs.

**Fig. S5**

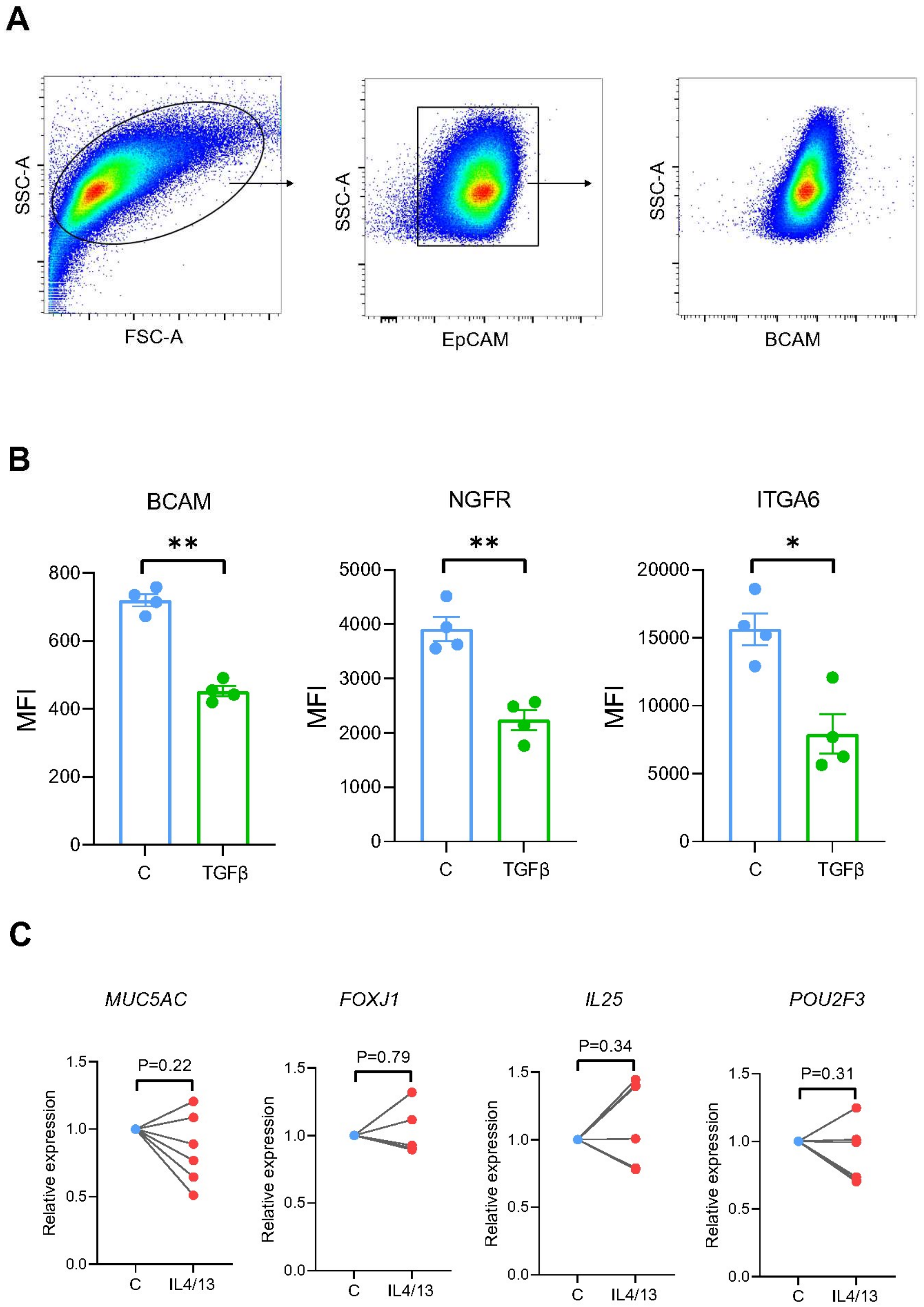
A
SSC-A
FSC-A
SSC-A
EpCAM
SSC-A
BCAM
B
BCAM
**
MFI
800
600
400
200
0
C
TGFβ
NGFR
**
MFI
5000
4000
3000
2000
1000
0
C
TGFβ
ITGA6
*
MFI
20000
15000
10000
5000
0
C
TGFβ
C
MUC5AC
P=0.22
Relative expression
1.5
1.0
0.5
0.0
C
IL4/13
FOXJ1
P=0.79
Relative expression
1.5
1.0
0.5
0.0
C
IL4/13
IL25
P=0.34
Relative expression
1.5
1.0
0.5
0.0
C
IL4/13
POU2F3
P=0.31
Relative expression
1.5
1.0
0.5
0.0
C
IL4/13

**Fig. S5: Analysis of BC surface markers in passaged and stimulated BCs.**
(**A**) Flow cytometry demonstrating BCAM expression in passaged human sinonasal BC cultures. (**B**) MFI of the indicated cell surface markers on passaged human sinonasal BCs treated with or without TGFβ (10 ng/ml) for 48 h. Data are shown as mean ± SEM (*p <0.05, ** p <0.01; paired two-tailed t test). (**C**) qPCR for the indicated genes in human sinonasal BCs treated with and without IL-4/13 (10 ng/ml) for 48 h. Data are shown as mean ± SEM (*p<0.05, ** p<0.01; paired two-tailed t test).
